# Mesoderm-Derived PDGFRA^+^ Cells Regulate the Emergence of Hematopoietic Stem Cells in the Dorsal Aorta

**DOI:** 10.1101/2021.08.08.455592

**Authors:** Vashe Chandrakanthan, Prunella Rorimpandey, Fabio Zanini, Diego Chacon, Young Chan Kang, Kathy Knezevic, Yizhou Huang, Qiao Qiao, Rema A. Oliver, Ashwin Unnikrishnan, Daniel R. Carter, Brendan Lee, Chris Brownlee, Carl Power, Simon Mendez-Ferrer, Grigori Enikolopov, William Walsh, Berthold Göttgens, Samir Taoudi, Dominik Beck, John E. Pimanda

## Abstract

Mouse hematopoietic stem cells (HSCs) first emerge at embryonic day 10.5 (E10.5) on the ventral surface of the dorsal aorta, by endothelial-to-hematopoietic transition (EHT). We investigated whether cells with mesenchymal stem cell-like activity, which provide an essential niche for long-term HSCs (LT-HSCs) in the bone marrow, reside in the aorta- gonad-mesonephros (AGM) and contribute to the structural development of the dorsal aorta and EHT. Using transgenic mice, we demonstrate a lineage hierarchy for AGM stromal cells and traced the E10.5/E11.5 aortic endothelium and HSCs to mesoderm derived (*Mesp1*) PDGFRA^+^ stromal cells (*Mesp1*^der^ PSCs). *Mesp1*^der^ PSCs dominate the sub-endothelial and ventral stroma in the E10.5–E11.5 AGM but by E13.5 were replaced by neural crest (*Wnt1*) derived PDGFRA^+^ stromal cells (*Wnt1*^der^ PSCs). Co-aggregating non-hemogenic embryonic and adult endothelial cells with *Mesp1*^der^ PSCs but not with *Wnt1*^der^ PSCs resulted in activation of a hematopoietic transcriptional program in endothelial cells accompanied by EHT and generation of LT-HSCs. Dose-dependent inhibition of PDGFRA signalling or BMP, WNT, NOTCH signalling interrupted this reprogramming event. This partnership between endothelial cells and AGM *Mesp1*^der^ PSCs could potentially be harnessed to manufacture LT-HSCs from endothelium.

## INTRODUCTION

Hematopoietic stem cells (HSCs) have extensive self-renewal capacity and are the source of daughter cells that proliferate, mature, and develop into blood cells of all types ^1^. As such, understanding the rules that govern HSC emergence, proliferation, and maturation is important to reproduce these phenomena *in vitro* ^2^. Indeed, advances in knowledge of embryonic hematopoiesis have informed methods that have been used to produce HSC-like cells *in vitro* ^3–5^.

The hematopoietic system in the embryo develops in successive waves ^6^. The first blood progenitors to emerge (from the extra-embryonic yolk sac) are primitive erythrocytes, followed by erythroid–myeloid progenitors ^7^. In mouse embryos the first HSCs appear mid-gestationally (E10.5) from hemogenic endothelial cells ^8–10^ lining the ventral surface of the dorsal aorta through endothelial-to-hematopoietic transition (EHT) ^11, 12^, in a region known as the aorta-gonad-mesonephros (AGM) ^13, 14^. These HSCs are amplified in the fetal liver ^13^ and the placenta ^15, 16^; they take up residence in the bone marrow, which will serve as the major adult site of hematopoiesis. Hemogenic endothelium is specified between E8.5 and E10.5 ^17^ and progress through pre-HSC stages to generate HSCs in the AGM between E10.5–E12.5 ^18^. HSC development in the AGM is influenced by NOTCH ^19^, WNT ^20^, BMP ^21, 22^, and other signals ^23, 24^ from surrounding cells ^25^. These signals facilitate hematopoiesis in part by regulating the expression of critical hematopoietic transcription factors, including components of the FLI1, GATA2, and SCL transcriptional network, GFI1/GFI1B, and RUNX1^26–28^.

Mesenchymal stem cells (MSCs) and their progeny are important constituents of the niche that regulates the size of the HSC pool in adult bone marrow ^29, 30^. Although there are resident stromal cells in the AGM that support hematopoiesis ^31, 32^, little is known about their developmental origins, transcriptional and functional identity, and contributions to the generation of long-term repopulating HSCs (LT-HSCs). Unlocking these properties might give insight into improving current protocols aimed at the *in vitro* generation of LT-HSCs with a view to eventual therapeutic application.

In this study we used a library of reporter mice to systematically investigate the stromal cell fraction in the AGM. In doing so, we identified *Mesp1*-derived PDGFRA^+^ stromal cells (PSCs) as precursors of the aortic endothelium and regulators of LT-HSC emergence from the endothelium. *Mesp1*-derived PSCs (*Mesp1^der^* PSCs) are a transient cell population, distributed in the ventral stroma during AGM hematopoiesis and replaced by *Wnt1*-derived PSCs (*Wnt1^der^* PSCs) as LT-HSCs cease to emerge. Co-aggregation of *Mesp1^der^* PSCs with non-hemogenic embryonic and adult endothelial cells resulted in the generation of endothelial-derived LT-HSCs, and this process was shown to be dependent on a PDGF-AA/PDGFR signaling axis and WNT, BMP and NOTCH signaling activity.

## RESULTS

### The E11.5 AGM has resident long- and short-term repopulating CFU-Fs that can be discriminated by the presence of PDGFRA and *Nestin* expression

Although the existence of stromal cells in the AGM that support hematopoiesis is known, and AGM-derived stromal cell lines have proven to be a powerful tool for the identification of environmental HSC regulators ^21, 31, 33^, we lack knowledge of the characteristics of these cells and their influence on EHT. It has previously been reported that all bone marrow MSCs (BM-MSCs) in *Nestin*-GFP transgenic mice, where expression of green fluorescent protein (GFP) is regulated by *Nes* ^34^, were GFP^+^, and that ablation of these BM-MSCs resulted in significant loss of long-term repopulating hematopoietic stem cells (LT-HSCs) in 12–16-week-old mice ^30^. We therefore used *Nestin*-GFP transgenic mice to investigate stromal cell populations in the AGM (Fig. 1A).

**Fig 1.**
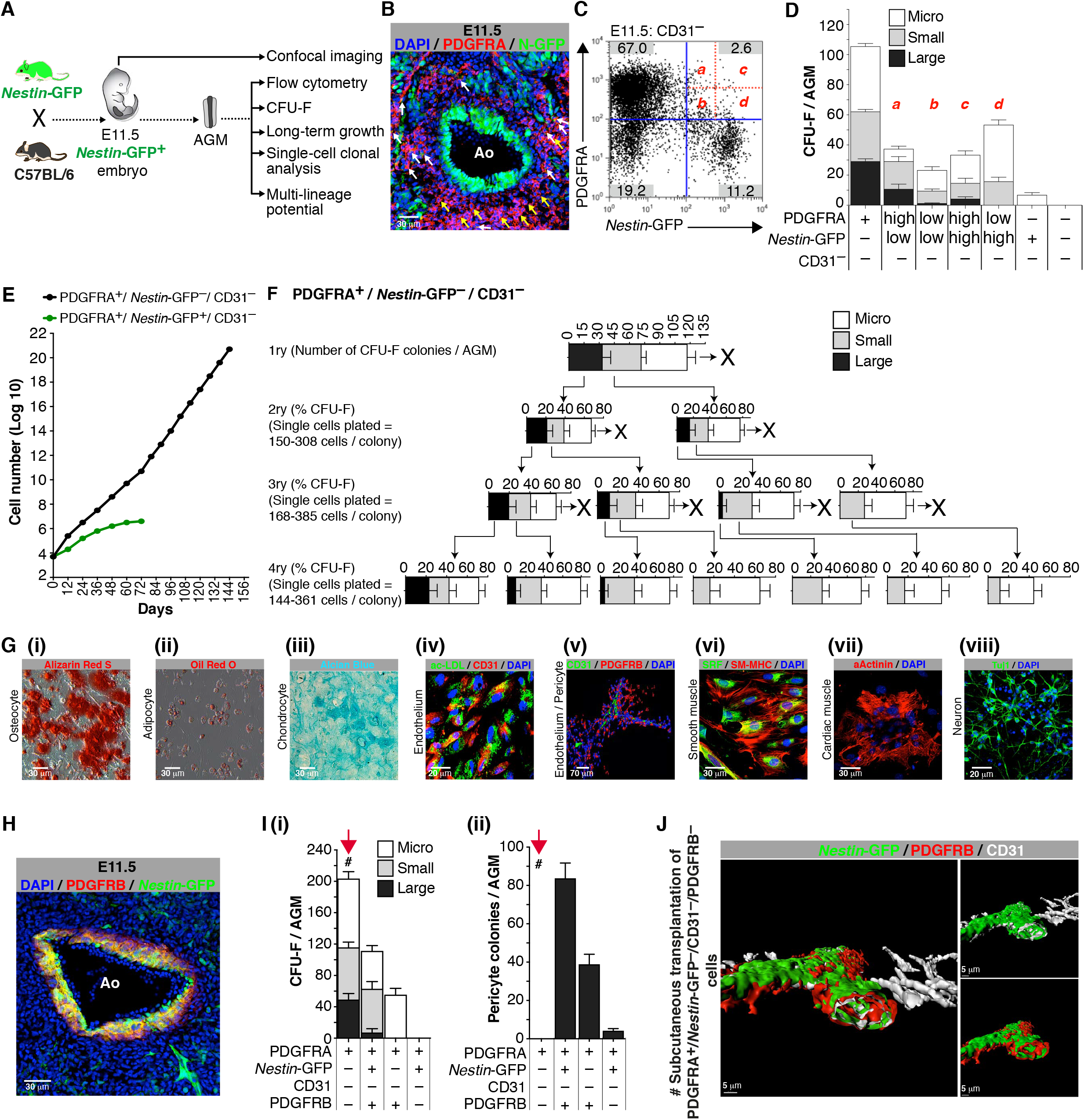
The E11.5 AGM has resident long- and short-term repopulating CFU-Fs that can be discriminated by expression of PDGFRA and *Nestin*-GFP. (A) Schematic outline of experiments performed using E11.5 *Nestin*-GFP^+^ embryos. (B) Confocal image of an E11.5 *Nestin*-GFP^+^ dorsal aorta stained for PDGFRA. (C) Flow cytometry analysis of E11.5 *Nestin*-GFP^+^ AGM (n=3) showing that 1:5.3 CD31^−^*Nestin*-GFP^+^ stromal cells are also PDGFRA^+^. (D) CFU-F potential of E11.5 CD31^−^; *Nestin*-GFP^+^ AGM (n=5) cells, sorted based on CD31, PDGFRA, and *Nestin*-GFP expression. PDGFRA^+^; *Nestin*-GFP^+^ cells were further fractionated into high and low positive sub-populations (a, b, c, d). (E) Long-term growth of E11.5 *Nestin*-GFP^+^ AGM-derived CFU-Fs based on CD31, *Nestin*-GFP, and PDGFRA expression (average values from n=3). (F) Single cell clonal analysis of CD31^−^; PDGFRA^+^; *Nestin*-GFP^−^ CFU-Fs. (G) *In vitro* differentiation of CD31^−^; PDGFRA^+^; *Nestin*-GFP^−^ cells (n=3). (H) Confocal image of an E11.5 *Nestin*-GFP^+^ dorsal aorta showing that a subset of PDGFRB^+^ cells co-express *Nestin*-GFP (n=3). PDGFRB^+^; *Nestin*-GFP^+^: yellow, PDGFRB^+^; *Nestin*-GFP^−^: red. (I) (i) CFU-Fs in FAC-sorted fractions from E11.5 *Nestin*-GFP^+^ AGMs (n=4). (ii) Pericyte colony-forming potential in FAC-sorted fractions from E11.5 *Nestin*-GFP^+^ AGMs (n=7). (J) Z-stack reconstruction of confocal images showing vessel-like structures lined by *Nestin*-GFP^+^ endothelial cells with surrounding PDGFRB^+^ pericytes (inset shows the lumen formed in the vessel). These images were taken from tissues harvested at 3 weeks after subcutaneous transplantation of a Matrigel plug loaded with PDGFRA^+^ *Nestin*-GFP^−^CD31^−^ PDGFRB^−^ FAC-sorted cells from E11.5 *Nestin*-GFP^+^ AGMs (see # in I (i) and (ii); red arrows). GFP: green fluorescent protein; AGM: aorta gonad mesonephros; Ao: aortic lumen; DAPI: 4’,6-diamidino-2-phenylindole dihydrochloride; PDGFRA: platelet-derived growth factor receptor alpha; PDGFRB: platelet-derived growth factor receptor beta; CFU-F: colony-forming unit–fibroblast; colony sizes: micro colonies (<2 mm, 2–24 cells), small colonies (2–4 mm, >25 cells), and large colonies (>4 mm, >100 cells); ** p<0.01; unpaired two-tailed t-test. Data represent mean ± s.e.m.

Confocal imaging of the E11.5 AGM of these mice showed that aortic endothelial, sub-endothelial, and blood cells adjacent to the aortic endothelium were *Nestin*-GFP^+^ (Extended Data Fig. 1A). Both *Nestin*-GFP^+^ and *Nestin*-GFP^−^ stromal cell fractions in the E11.5 AGM were found to express platelet-derived growth factor receptor A (PDGFRA), a tyrosine kinase receptor expressed on the surface of MSCs ^35^ (Fig. 1B and 1C) and on early embryonic mesodermal cells that contribute to hemogenic endothelium and hematopoietic cells ^36^. These PDGFRA^+^ cells (*Nestin*-GFP^−^, yellow arrows; and *Nestin*-GFP^+^, white arrows) were distributed deeper in the aortic parenchyma and surrounded the PDGFRA^−^/ *Nestin*-GFP^+^ cells, which were concentrated more toward the aortic lumen (Fig. 1B).

To explore transitional functional properties of PDGFRA and *Nestin*-GFP expressing and non-expressing cells in the E11.5 AGM, we used an *in vitro* colony-forming unit– fibroblast (CFU-F) assay ^37^. Freshly isolated FAC-sorted PDGFRA^+^/ *Nestin*-GFP^−^ and PDGFRA^+^/ *Nestin*-GFP^+^ populations produced CFU-Fs of varying sizes ^38^ and had mesenchymal cell morphology (Fig. 1D and Extended Data Fig. 1B). Although both PDGFRA^+^/ *Nestin*-GFP^−^ and PDGFRA^+^/ *Nestin*-GFP^+^ cells produced large CFU-F colonies (Fig. 1D), their number was lower in the latter and proportionate to the intensity of PDGFRA and *Nestin*-GFP expression. Only PDGFRA^+^/ *Nestin*-GFP^−^ cells showed long-term re-plating capacity (Fig. 1E).

Furthermore, serial re-plating of single cells from PDGFRA^+^/ *Nestin*-GFP^−^ large CFU-F colonies produced consistent numbers of large colonies of CFU-Fs (Fig. 1F), and these cells could be differentiated *in vitro* into mesodermal and ectodermal derivatives (Fig. 1G (i)-(viii) and movie 1; osteocytes, adipocytes, chondrocytes, endothelium, smooth muscle, beating cardiomyocytes—movie 1—and neurons). By contrast, single cells from PDGFRA^+^/ *Nestin*-GFP^+^ large CFU-F colonies showed limited capacity to generate large colonies of CFU-Fs through serial single cell re-plating (Extended Data Fig. 1C); they could only be differentiated into adipocytes, endothelium, and smooth muscle (Extended Data Fig. 1D-E). The differentiation potential observed in bulk PDGFRA^+^/*Nestin*-GFP^−^ and PDGFRA^+^/*Nestin*-GFP^+^ cells was best reflected in cells that formed large CFU-F colonies (Extended Data Fig. 1F). Taken together, these data show that PDGFRA marks all CFU-Fs in the E11.5 AGM, with *Nestin* expression marking a sub-population of PDGFRA^+^ cells with limited potential for CFU-F activity and differentiation.

Pericytes are characterized by PDGFRB expression ^39, 40^ and were distributed concentrically in the sub-endothelium of the E11.5 dorsal aorta (Extended Data Fig. 1H). To investigate the relationship between *Nestin*-GFP^+^ cells and pericytes, we fractionated cell populations (FAC-sorted) based on PDGFRA, *Nestin*-GFP, CD31, and PDGFRB from E11.5 AGMs of *Nestin*-GFP transgenic mice (Extended Data Fig. 1G) and performed assays for formation of CFU-F and pericyte colonies and long-term re-plating assay (Fig. 1I (i)-(ii) and Extended Data Fig. 1H). Among CD31^−^/PDGFRA^+^ cells, PDGFRB co-expression was proportionately higher in *Nestin*-GFP^+^ than in *Nestin*-GFP^−^ cells (Extended Data Fig. 1G). While the latter showed the highest large-CFU-F colony and long-term re-plating potential, unlike the former cells they lacked potential to form pericyte colonies (Fig. 1I (i)-(ii) and Extended Data Fig. 1H). Interestingly, CFU-F potential in the *Nestin*-GFP^+^ fraction was exclusively within the PDGFRB^+^ sub-fraction (Fig. 1I (i)). We further assessed the contribution of different PDGFRA^+^ fractions (Fig. 1I) toward *in vivo* morphological and functional vascular contents. We purified PDGFRA^+^ fractions using flow cytometry (Fig. 1I and Extended Data Fig. 1G), mixed those cells with matrigel, and transplanted them subcutaneously into C57BL/6 mice. Only purified CD31^−^/PDGFRA^+^/*Nestin*-GFP^−^/PDGFRB^−^ CFU-Fs formed branching vessel-like structures (Fig. 1J and Movie 2)—the luminal surfaces of which were lined with *Nestin*-GFP^+^/CD31^+^ endothelial cells and enveloped by PDGFRB^+^ pericytes.

In summary, at E11.5 the mouse AGM has a population of resident PDGFRA^+^ stromal cells distributed around the lumen of the developing aorta. Expression of *Nestin*-GFP^+^, PDGFRB, or CD31 in PDGFRA^+^ cells was found to coincide with progressively restricted potential for proliferation, differentiation, and contribution to vascular components such as endothelium and pericytes.

### E9.5 PDGFRA^+^ cells contribute to hemogenic endothelium and LT-HSCs

If PDGFRA^+^ stromal cells are a reservoir for endothelial and sub-endothelial cells in the developing aorta, they could potentially also contribute to long-term repopulating HSCs that emerge at E11.5. To explore whether cells expressing PDGFRA and CD31 proteins in the AGM were early and late constituents of a differentiation continuum, we evaluated the distribution of CD31 in the E8.5, E9.5, E10.5 and E11.5 AGMs of *Pdgfra*-nGFP knock-in mice (that is, mice whose *Pdgfra*-expressing cells retain GFP in their nuclei) ^41^ (Fig. 2A (i)). *Pdgfra*-nGFP cells are in proximity with endothelial cells of the paired dorsal aorta at early embryonic timepoints (Fig. 2A (ii)). At E11.5 (Fig. 2A (iii)), cells furthest from the aortic lumen showed robust *Pdgfra*-nGFP^high^ expression but no NESTIN (NES) protein (layer I). Cells co-expressing both *Pdgfra*-nGFP^high^ and NES (layer II) were interspersed between these cells (layer I) and cells that were *Pdgfra*-nGFP^low^ but NES^+^ (layer III), which also co-expressed the smooth muscle marker aSMA (Extended Data Fig. 2A). Endothelial cells lining the aortic lumen (layer V) were *Pdgfra*-nGFP*^−^* and expressed CD31 but little or no NES (in contrast to the longer-lasting GFP in N-GFP transgenic mice; Fig. 1A). A few cells were NES^+^ and CD31^+^ but low in *Pdgfra*-nGFP (layer IV). It is salient that *Pdgfra*-nGFP^+^ and PDGFRA^+^ cells in the E11.5 AGM were comparable in their CFU-F potential and long-term growth potential (Extended Data Fig. 2B and 2C (i)-(iii)).

**Fig 2.**
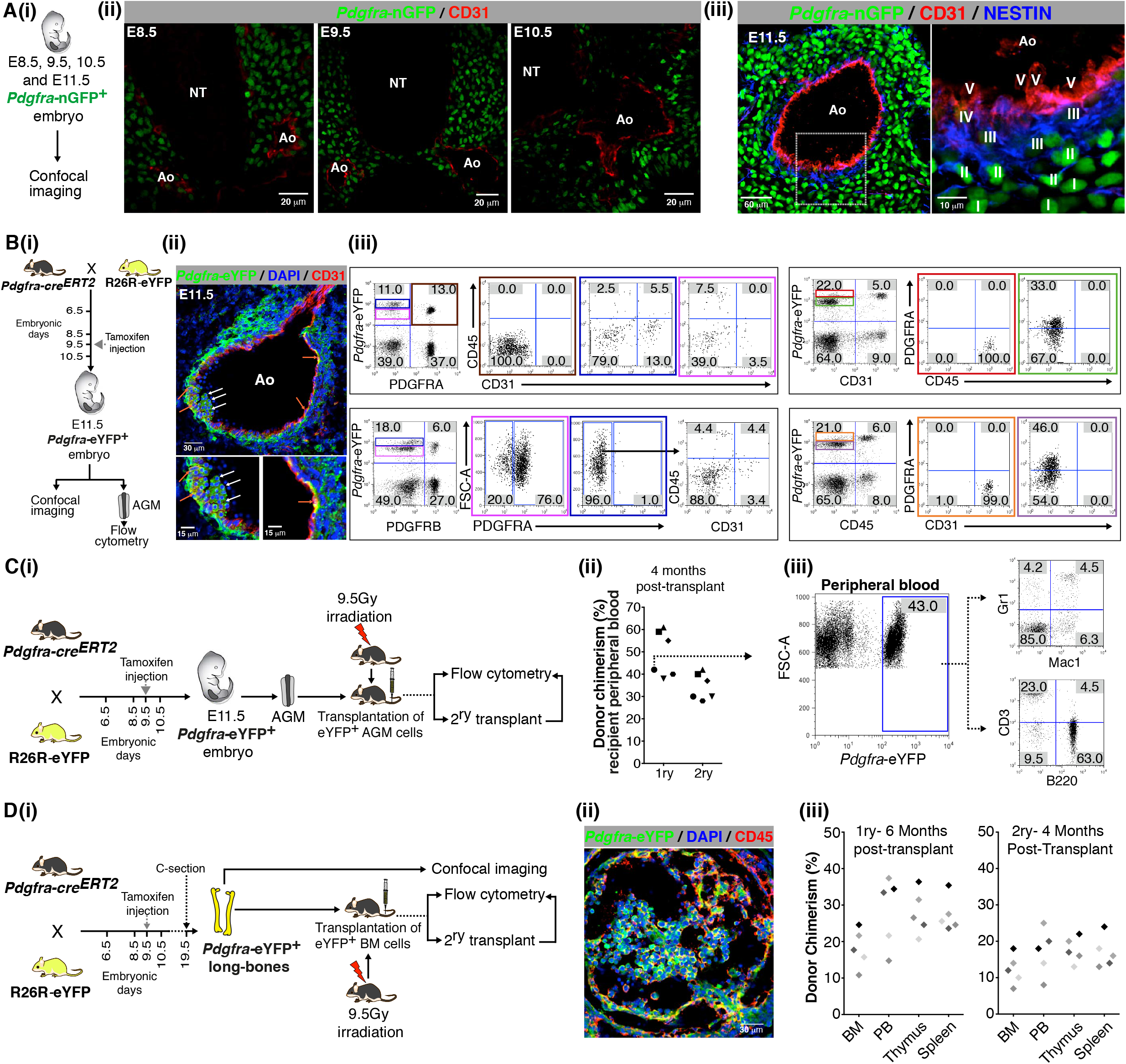
E9.5 PDGFRA^+^ cells contribute to hemogenic endothelium and LT-HSCs. (A) (i) Schematic outline of experiments performed using E8.5, E9.5, E10.5 and E11.5 *Pdgfra*-nGFP embryos. (ii) Confocal images of E.8.5, E9.5 and E10.5 *Pdgfra*-nGFP embryos showing the distribution of PDGFRA expressing cells in relation to the developing aorta. (iii) Spatial distribution of *Pdgfra*-nGFP, NESTIN, and CD31 expressing cells in a *Pdgfra*-nGFP E11.5 AGM. Cell populations are marked I to V: I) *Pdgfra*-nGFP^high^, NESTIN^−^, CD31^−^; II) *Pdgfra*-nGFP^high^, NESTIN^+^, CD31^−^; III) *Pdgfra*-nGFP^low^, NESTIN^+^, CD31^−^; IV) *Pdgfra*-nGFP^low^, NESTIN^+^, CD31^+^; V) *Pdgfra*-nGFP^−^, NESTIN^−^, CD31^+^. (B) (i) Schematic outline of lineage-tracing experiments using *Pdgfra-cre^ERT^*^2^ *R26R*-eYFP embryos. (ii) Confocal image of an E11.5 *Pdgfra-cre^ERT^*^2^ *R26R*-eYFP AGM following *cre*-activation at E9.5, showing eYFP^+^ blood cells (white arrows) and endothelium (orange arrows). (iii) Contribution of donor eYFP^+^ cells to PDGFRA, pericytes (PDGFRB), endothelium (CD31), and blood cells (CD45) in the E11.5 *Pdgfra-cre^ERT^*^2^ *R26R*-eYFP AGM following *cre*-activation at E9.5. (C) (i) Schematic outline of lineage-tracing experiments using *Pdgfra-cre^ERT2^ R26R*-eYFP embryos. (ii) Contribution of donor eYFP^+^ cells to peripheral blood in primary and secondary transplants. (iii) Flow cytometry analysis of donor eYFP^+^ cells to peripheral blood in primary transplant. (D) (i) Schematic outline of lineage-tracing experiments using *Pdgfra-cre^ERT2^ R26R*-eYFP embryos. (ii) Confocal image of a *Pdgfra-cre^ERT2^ R26R*-eYFP neonatal long-bone section following *cre*-activation at E9.5, showing eYFP^+^ (green) CD45^+^ (red) blood cells in the bone marrow. (iii) Contribution of donor eYFP^+^ cells to peripheral blood, bone marrow, thymus, and spleen in primary and secondary transplants. GFP: green fluorescent protein; eYFP; enhanced yellow fluorescent protein; Ao: aortic lumen; NT: neural tube; AGM: aorta gonad mesonephros; DAPI: 4’,6-diamidino-2-phenylindole dihydrochloride; FSC-A: forward scatter area; PB: peripheral blood; BM: bone marrow.

To formally establish a lineage relationship between PDGFRA^+^ cells and their progeny, we crossed *Pdgfra-cre^ERT2^* ^42^ with R26R-eYFP ^43^ mice to generate *Pdgfra-cre^ERT2^*/R26R-eYFP compound transgenic embryos (Fig. 2B (i)) and induced *cre*-recombination at E9.5 by delivering single injections of tamoxifen to pregnant mothers and harvesting embryos at E11.5. CD31^+^ endothelial cells in the E9.5 AGM do not express *Pdgfra* (Extended Data Fig. 2D (i)-(ii)) ^44^. There was efficient recombination with 6.4% of limb bud cells expressing eYFP following a single injection of tamoxifen at E9.5 (Extended Data Fig. 2E). The *Pdgfra-cre^ERT^*^2^/R26R-eYFP recombination at E9.5 resulted in eYFP^+^ aortic endothelial, sub-endothelial, and blood cells in the E11.5 AGM (Fig. 2B (ii)-(iii)) with only a minority of eYFP^+^ cells still expressing PDGFRA protein (Fig. 2B (iii) and Extended Data Fig. 2F). eYFP^+^/PDGFRA^+^ cells were CD31 and CD45 negative and had lower eYFP fluorescence than CD31 or CD45 positive cells (Fig. 2B). There were no eYFP^+^ CD31^+^ endothelial cells in the E11.5 yolk sac, placenta, or umbilical and vitelline vessels (Extended Data Fig. 2G).

To evaluate whether these eYFP^+^ cells included LT-HSCs, we again induced cre-recombination in *Pdgfra-cre^ERT2^*/R26R-eYFP compound transgenic embryos at E9.5, harvested E11.5 embryos, and performed transplantation assays with eYFP^+^ AGM cells (Fig. 2C (i)). These cells were able to reconstitute hematopoiesis in lethally irradiated mice following primary and secondary transplantation and contributed to multiple blood lineages (Fig. 2C (ii)-(iii) and Extended Data Fig. 2H). To establish whether *Pdgfra*-eYFP^+^ cells populate the bone marrow, *Pdgfra-cre^ERT2^*/R26R-eYFP compound transgenic embryos were matured to term following induction of recombination at E9.5 and delivered by caesarean section (owing to difficulties in parturition) (Fig. 2D (i)). In *Pdgfra-cre^ERT2^/*R26R-eYFP compound neonatal mice, eYFP^+^/CD45^+^ blood cells were present in the BM (Fig. 2D (ii)). These cells were able to reconstitute hematopoiesis in lethally irradiated mice following primary and secondary transplantation (Fig. 2D (iii)) and contributed to multiple blood lineages (Extended Data Fig. 2I).

Given the contributions of E9.5 PDGFRA^+^ cells to structures of the aorta and blood cells that arise therein at E11.5, we predicted that ablating these cells and their progeny would have a profoundly deleterious impact on the developing aorta and hematopoiesis. To explore this, we crossed *Pdgfra*-*cre^ERT2^* mice ^42^ with inducible diphtheria toxin receptor (iDTR) mice ^45^ to generate *Pdgfra-cre^ERT2^*/iDTR embryos (Extended Data Fig. 2J). We conditionally induced expression of diphtheria toxin receptor in E9.5 PDGFRA^+^ cells by treatment with tamoxifen, followed by ablation of these cells using diphtheria toxin (DT) at E10.5 in *Pdgfra-cre^ERT2^*/iDTR embryos (Extended Data Fig. 2J (i)). We then studied the resulting impact on AGM architecture at E11.5. In whole-mount and tissue sections of compound transgenic embryos, there was severe disruption of normal dorsal aorta development (Extended Data Fig. 2J (ii)). In these embryos there was concomitant reduction in numbers of various cell types: endothelial (CD31^+^); blood (SCA1^+^; CD45^+^); perivascular (PDGFRB^+^); CFU-Fs (PDGFRA^+^) (Extended Data Fig. 2J (iii)), as well as blood progenitors (Extended Data Fig. 2J (iv)) and CFU-Fs (Extended Data Fig. 2J (v)). These data indicate that the absence of PDGFRA expressing cells in the developing embryo should have a profoundly deleterious impact on AGM hematopoiesis. Mice carrying a targeted null mutation of *Pdgfra* show early embryonic lethality ^46^ and PDGFRA signalling has previously been reported to be essential for establishing a microenvironment that supports definitive hematopoiesis ^47^. To directly test whether LT-HSCs were generated in the absence of *Pdgfra*, we crossed tdTomato/Rosa26; *Pdgfra*-nGFP knock-in heterozygote mice to generate *Pdgfra* KI/KI (null) and KI/+ (heterozygote) embryos with ubiquitous tdTomato expression and performed CFU-C and transplantation assays with individual E10.5 and E11.5 AGMs from GFP^+^ KI embryos with retrospective genotyping of yolk sacs (Extended Data Fig. 2K (i)). Consistent with our expectations, *Pdgfra* null E10.5 and E11.5 AGMs produced significantly fewer CFU-Cs and no LT-HSCs (Extended Data Fig. 2K (ii) and (iii)).

Taken together, these data show that E11.5 AGM LT-HSCs, aortic endothelium and surrounding accessory cells were in part derived from PDGFRA^+^ precursors in the E9.5 AGM and that PDGFRA expressing cells were required for the generation of LT-HSCs.

### Transient and distinct waves of PSCs serially populate the AGM

MSCs in the trunks of E14.5 embryos have previously been reported to be derived from Sox1^+^ neuroepithelium, in part through a neural crest intermediate ^35^. But even at this stage of development it was evident that PDGFRA^+^ MSC progenitors that were derived from neither the neuroepithelium nor the neural crest accounted for most MSC populations in postnatal bone marrow ^35^.

To investigate the source of AGM CFU-Fs, we first crossed R26R-eYFP mice with *Mesp1-cre* (mesoderm) mice ^48^ and harvested embryos for confocal imaging, and AGM flow cytometry and CFU-F assays (Fig. 3A (i)). At E11.5, CD31^+^ aortic endothelial cells were *Mesp1*-eYFP^+^ and were surrounded by a rim of *Mesp1*-eYFP^+^/CD31^−^ sub-endothelial cells (Fig. 3A (ii; left)). Sub-endothelial cells expressing the smooth muscle marker Calponin were also *Mesp1*-eYFP^+^ (Fig. 3A (ii; right)). A survey of E8.5, E9.5 and E10.5 AGM in *Mesp1*-eYFP^+^ embryos showed that *Mesp1* derived stromal cells also contributed to the aortic endothelium even at these early timepoints (Extended Data Fig. 3A). These data collectively show that at the time of HSC emergence, sub-endothelial stromal cells were mesodermal derivatives.

**Fig 3.**
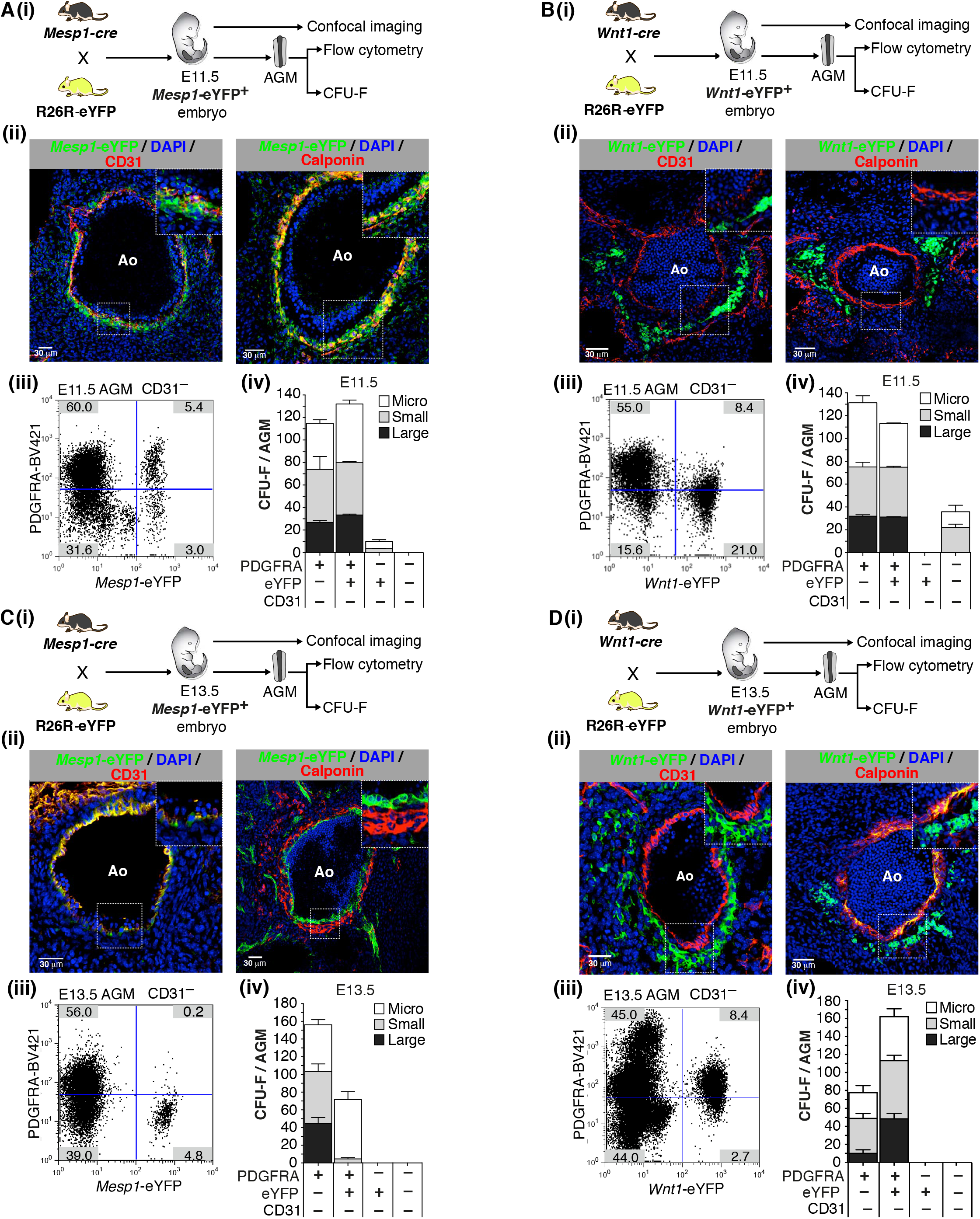
Developmental origins of AGM endothelium and CFU-Fs. (A) (i) Schematic outlining the genetic cross used to harvest *Mesp1-cre R26R*-eYFP (*Mesp1*-eYFP^+^) embryos at E11.5. (ii) Confocal images of E11.5 *Mesp1*-eYFP AGM showing contribution of *Mesp1*-derived cells to the endothelium (left; CD31) and smooth muscle (right; Calponin). (iii) Flow cytometry showing percentage of CD31^−^; *Mesp1*-eYFP^+^ PDGFRA^+^ cells in AGMs at E11.5. (iv) CFU-Fs in cell fractions sorted from *Mesp1*-eYFP^+^ AGMs (n=7) at E11.5. (B) (i) Schematic outlining the genetic cross used to harvest *Wnt1*-eYFP embryos at E11.5. (ii) Confocal images of E11.5 *Wnt1*-eYFP AGM showing absence of contribution to endothelium (left; CD31) or to sub-endothelial smooth muscle (right; Calponin) or to sub-endothelial stroma (left and right). (iii) Flow cytometry showing percentage of CD31^−^; *Wnt1*-eYFP^+^ PDGFRA^+^ in AGMs at E11.5. (iv) CFU-Fs in cell fractions sorted from *Wnt1*-eYFP^+^ AGMs (n=5) at E11.5. (C) (i) Schematic outlining the genetic cross used to harvest *Mesp1-cre R26R*-eYFP (*Mesp1*-eYFP^+^) embryos at E13.5. (ii) Confocal images of E13.5 *Mesp1*-eYFP AGM showing contributions of *Mesp1*-derived cells to the endothelium (left; CD31) but not to smooth muscle (right; Calponin). (iii) Flow cytometry showing percentage of CD31^−^; *Mesp1*-eYFP^+^ PDGFRA^+^ cells in AGMs at E13.5. (iv) CFU-Fs in cell fractions sorted from *Mesp1*-eYFP^+^ AGMs (n=7) at E13.5. (D) (i) Schematic outlining the genetic cross used to harvest *Wnt1*-eYFP embryos at E13.5. (ii) Confocal images of E13.5 *Wnt1*-eYFP AGM showing absence of contributions of *Wnt1* derived cells to the endothelium (left; CD31), but contributions to smooth muscle (right; Calponin) and to sub-endothelial stroma (left and right). (iii) Flow cytometry showing percentage of CD31^−^; *Wnt1*-eYFP^+^ PDGFRA^+^ in AGMs at E13.5. (iv) CFU-Fs in cell fractions sorted from *Wnt1*-eYFP^+^ AGMs (n=5) at E13.5 (right). Ao: aortic lumen; FAC flow activated cell; DAPI: 4’,6-diamidino-2-phenylindole dihydrochloride; BV421-Brilliant Violet 421; CFU-F: colony-forming unit–fibroblast. Colony sizes: Micro colonies (<2 mm, 2–24 cells), small colonies (2–4 mm, >25 cells) and large colonies (>4 mm, >100 cells). Data represent mean ± s.e.m.

Approximately, 2:3 *Mesp1*-eYFP^+^ cells were PDGFRA^+^ (Fig. 3A (iii)). Indeed, in E11.5 *Pdgfra*-nGFP^+^/*Mesp1*-DsRed double transgenic embryos, the AGM had GFP^+^ stromal cells, endothelial cells, and blood clusters that were also DsRed^+^ (Extended Data Fig. 3B). However, in both E11.5 and E10.5 embryos, *Mesp1*-eYFP^+^ and *Mesp1*-eYFP^−^ PDGFRA^+^ cells generated comparable numbers of CFU-Fs (Fig. 3A (iv); Extended Data Fig. 3C). PDGFRA^−^ cells on the other hand had limited CFU-F capacity at both time points and formed no large colonies. Taken together these data show that the aortic endothelium, sub-endothelium, and a proportion of CFU-Fs in the E11.5 AGM were derived from *Mesp1*^+^ cells but that a comparable number of CFU-Fs were not.

To explore whether the *Mesp1*-eYFP^−^ cells were derived from *Wnt1^+^* cells, we next crossed R26R-eYFP mice with *Wnt1-cre* (neural crest) mice ^49^ and harvested embryos for confocal imaging, and AGM flow cytometry and CFU-F assays (Fig. 3B (i)). In contrast to *Mesp1*-eYFP^+^ cells at a corresponding embryonic time-points (E8.5-E11.5), *Wnt1*-eYFP^+^ cells did not contribute to the endothelium or sub-endothelium and were located deeper in the AGM stroma (E11.5; Fig. 3B (ii)) or distant to the ventral surface of the dorsal aorta (E8.5-E10.5; Extended Data Fig. 3D). However, 1:3 *Wnt1*-eYFP^+^ cells were PDGFRA^+^ (Fig. 3B (iii)) and these cells formed CFU-Fs with comparable efficiency to *Mesp1*-eYFP^+^/PDGFRA^+^ cells (compare Fig. 3B (iv) with Fig. 3A (iv) and F Extended Data Fig. 3E with Extended Data Fig. 3C) in both E11.5 and E10.5 embryos.

Unlike MSCs in the E14.5 embryonic trunk, which were reported to be derived from Sox1^+^ neuroepithelium ^35^, *Sox1*-eYFP^+^/PDGFRA^+^ cells were very rare in the E11.5 AGM (2%) and did not contribute to large CFU-Fs (Extended Data Fig. 3F (i)-(iii)). Therefore, the E11.5 AGM has at least two populations of PDGFRA^+^ CFU-Fs that have different lineage ancestries (*Mesp1*^der^ and *Wnt1*^der^) and occupy distinct anatomical locations with respect to the hemogenic endothelium.

In contrast to the hemogenic E10.5/E11.5 AGM, the E13.5 AGM is no longer hemogenic ^18^. To evaluate whether CFU-F populations changed during this transition, we crossed R26R-eYFP mice with *Mesp1-cre* mice and harvested embryos for confocal imaging, and AGM flow cytometry and CFU-F assays at E13.5 (Fig. 3C (i)). Whereas the dorsal aorta was still lined by *Mesp1*-eYFP^+^/CD31^+^ endothelial cells (Fig. 3C (ii; left)), Calponin^+^ sub-endothelial cells were *Mesp1*-eYFP^−^ (Fig. 3C (ii; right)). Furthermore, *Mesp1*-eYFP^+^/PDGFRA^+^ stromal cells, which were relatively abundant at E11.5 (5.4%; Fig. 3A (iii)) were rare at E13.5 (0.2%; Fig. 3C (iii)). Large-CFU-F potential in the E10.5 and E11.5 AGM was seen in both *Mesp1*-eYFP^+^ and eYFP^−^ PDGFRA^+^ fractions (Fig. 3A (iv)) and Extended Data Fig. 3C) but in the absence of *Mesp1*-eYFP^+^/PDGFRA^+^ cells at E13.5, they were derived exclusively from *Mesp1*-eYFP^−^/PDGFRA^+^ cells (Fig. 3C (iv)).

We then crossed R26R-eYFP mice with *Wnt1-cre* mice and harvested embryos at E13.5 for confocal imaging, and AGM flow cytometry and CFU-F assays (Fig. 3D (i)). As observed in E8.5-E11.5 embryos (Fig. 3B (ii; left) and Extended Data Fig. 3D), there was no evidence of *Wnt1*-eYFP^+^-derived endothelial cells at E13.5 (Fig. 3D (ii; left)) but the layer of *Mesp1*-eYFP^+^/CD31^−^/Calponin^+^ sub-endothelial cells that were evident at E11.5 had been replaced by *Wnt1*-eYFP^+^/CD31^−^/Calponin^+^ cells (Fig. 3D (ii; right)). There were equal proportions (8.4%) of *Wnt1-eYFP* PSCs at E13.5 as at E11.5 (compare Fig. 3D (iii) with 3B (iii)). In the absence of *Mesp1*-eYFP^+^ PSCs at E13.5, large CFU-F potential was seen largely in *Wnt1*-eYFP^+^ PSCs (Fig. 3D (iv)).

It is salient to note that *Mesp1* transcripts were absent in PDGFRA^+^ (CFU-F), PDGFRB^+^ (pericytes), or CD31^+^ (Endothelial) cells in the AGM at both E11.5 and E13.5 (Extended Data Fig. 4A). Therefore *Mesp1*-eYFP^+^ cells in the AGM at these time-points are Mesp1 derived cells that do not currently express *Mesp1*. Although *Wnt1* transcripts were absent in E13.5 cells, there was variable and low-level *Wnt1* expression at E11.5 in PDGFRA^+^ (CFU-F) but not PDGFRB^+^ (pericytes) or CD31^+^ (endothelial) cells (Extended Data Fig. 4B). As such, *Wnt1*-eYFP^+^ cells in the E13.5 AGM could be derivatives either of *Wnt1*-expressing E11.5 PDGFRA^+^ cells or from cells that expressed *Wnt1* at an earlier time point during development.

Taken together, these data show that at the time of HSC emergence at E11.5 (and E10.5), sub-endothelial stromal cells were mesodermal (i.e., *Mesp1*) derivatives. The loss of *Mesp1*-derived (*Mesp1^der^*) cells in the sub-endothelium, along with replacement by *Wnt1*-derived (*Wnt1^der^*) cells at E13.5, temporally coincides with the loss of EHT in the dorsal aorta. This raises the intriguing possibility that *Mesp1^der^* PSCs play an active role in EHT in contrast with *Wnt1^der^* PSCs.

### LT-HSC production can be induced in non-hemogenic endothelial cells by co-aggregating these cells with E11.5 mesoderm-derived PSCs

To determine whether there were EHT-promoting attributes in E10.5 or E11.5 *Mesp1^der^* PSCs that were absent in E11.5 or E13.5 *Wnt1*-derived progenitors, we performed co-aggregate cultures of FAC-sorted *Mesp1^der^*- or *Wnt1^der^* PSCs with endothelial cells from ubiquitous GFP^+^ (*UBC-gfp*/*BL6*) ^50^ mice. *Mesp1^der^*- or *Wnt1^der^*- PSCs were harvested from the AGMs of compound transgenic embryos generated by crossing *Mesp1*-*cre* or *Wnt1*-*cre* mice with *STOCK Tg(CAG-Bgeo-DSRed*MST)1Nagy/J* (Z/Red) reporter mice ^51^.

Endothelial cells (*UBC*-GFP^+^/PDGFRA^−^/PDGFRB^−^/CD31^+^/VE-Cad^+^/CD41^−^/CD45^−^) from E10.5, E11.5 and E13.5 AGM or 8–12-week-old adult mice (heart, aorta, and inferior vena cava (IVC)) were co-aggregated with stromal cells (PDGFRA^+^/PDGFRB^−^/CD31^−^/VE-Cad^−^/CD41^−^/CD45^−^) from E10.5 and E11.5 *Mesp1*-DSRed^+^ AGM (Fig. 4A (i)). There were no *Mesp1*^der^ PSCs in the E13.5 AGM (Fig. 3C (ii)) hence co-aggregates could not be performed with cells harvested from this embryonic time-point. Following 96 hours of culture, the co-aggregates were cryosectioned for confocal imaging (for evidence of CD45^+^ blood cells) or used for CFU-C and transplantation assays to establish progenitor and stem cell potential of emerging blood cells. Confocal microscopy showed colonies of GFP^+^/CD45^+^ cells in all cryosectioned endothelial/*Mesp1^der^*PSC co-aggregates. (Fig. 4A (ii); E13.5 AGM endothelium/11.5 *Mesp1^der^*PSCs). No DsRed^+^/CD45^+^ cells were found in any co-aggregate. Co-aggregation of both E10.5 and E11.5 *Mesp1*^der^ PSCs with E11.5 (hemogenic) or E13.5 (non-hemogenic) AGM or adult heart or aortic endothelium resulted in the emergence of CFU-Cs (Fig. 4A (iii)) and endothelial cell derived (*UBC*-GFP^+^) LT-HSCs with robust multi-lineage hematopoietic reconstitution (Fig. 4A (iv)-(v); Extended Data Fig. 5A-B). There were no PSC derived (DsRed^+^) hematopoietic cells in any co-aggregate transplant recipient (Extended Data Fig. 5A (ii)-(iv)). Transplantation of aggregates composed of endothelial cells or PSCs alone did not contribute to hematopoietic cells (Extended Data Fig. 5C). Co-aggregates of E10.5 and E11.5 *Mesp1*^der^ PSCs with IVC endothelium yielded CFU-Cs but their number was significantly lower than those from aortic endothelium with correspondingly low donor chimerism following transplantation (Fig. 4A (iv)-(v)).

**Fig 4.**
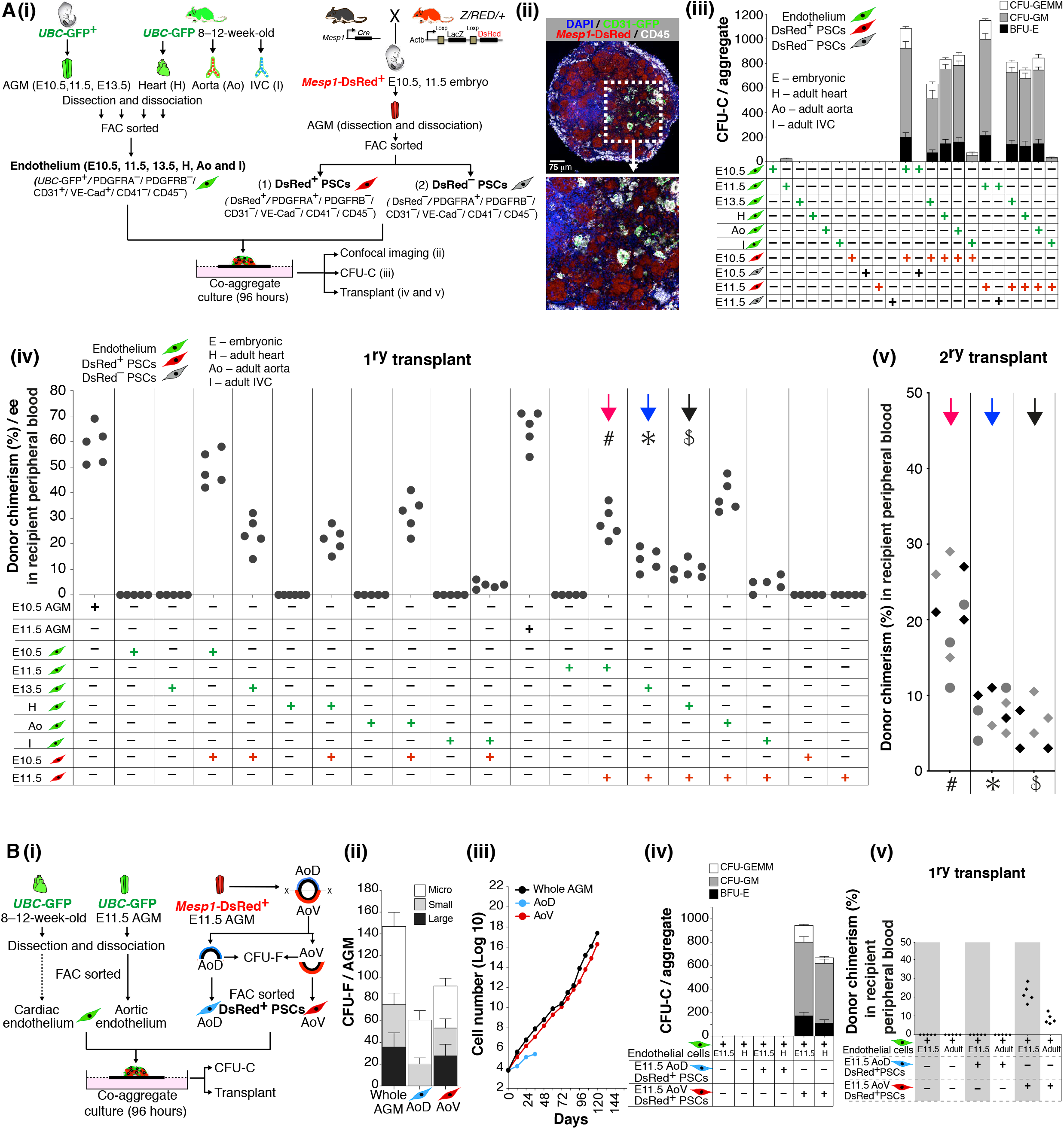
Co-aggregate cultures of endothelial cells with E11.5 *Mesp1*^der^ PSCs generate endothelium-derived LT-HSCs. (A) (i) Schematic outlining the process for harvesting cell types used in co-aggregate cultures. PSCs (150,000 cells) from E10.5 and E11.5 embryos were co-aggregated with endothelial cells from E10.5-, E11.5-, E13.5-AGM, Adult-heart, -aorta, -inferior vena cava (25,000 cells) and cultured for 96 hours. The PSCs were DsRed^+^ and DsRed^−^ (*Mesp1*-DsRed^+/–^/PDGFRA^+^/PDGFRB^−^/CD31^−^/VECad^−^/ CD41^−^/CD45^−^); the endothelial cells were GFP^+^ (*UBC*-GFP^+^/PDGFRA^−^/PDGFRB^−^/CD31^+^/VECad^+^/CD41^−^/CD45). (ii) Confocal images of cryosections of a co-aggregate of E11.5 *Mesp1*-DsRed^+^ PSCs and E13.5 endothelial cells at 96 hours, showing GFP^+^/CD45^+^ cells. (iii) CFU-C potential of embryonic and adult aortic endothelium co-aggregated with E10.5 or E11.5 *Mesp1*-DsRed^+^ or E11.5 *Mesp1*-DsRed^−^ PSCs. (iv) Percentage of GFP^+^ cells in peripheral blood of irradiated recipients six months after transplantation of co-aggregates (one co-aggregate for each adult irradiated recipient). (v) Percentage of GFP^+^ cells in peripheral bloods of irradiated recipients four months after bone marrow transplants from corresponding mice in (iv). (B) (i) Schematic outlining the process of harvesting endothelial cells from E11.5 AGM or adult heart and *Mesp1*^der^ PSCs from the dorsal or ventral halves of the E11.5 aorta for CFU-F assays and co-aggregate cultures. (ii) CFU-F activity of *Mesp1*^der^ PSCs from the whole AGM or dorsal or ventral halves of the E11.5 aorta. (iii) Long-term replating of *Mesp1*^der^ PSCs from the whole AGM or dorsal or ventral halves of the E11.5 aorta. (iv) CFU-C potential of embryonic or adult endothelium co-aggregated with E11.5 *Mesp1*^der^ PSCs from either the dorsal or ventral halves of the aorta. (v) Percentage of GFP^+^ cells in peripheral blood of irradiated recipients six months after transplantation of co-aggregates (one co-aggregate for each adult irradiated recipient). DAPI: 4’,6-diamidino-2-phenylindole dihydrochloride; CFU-C: colony-forming unit–culture; BFU-E: burst-forming unit–erythroid; CFU-GM; colony-forming unit–granulocyte/ macrophage; CFU-GEMM: colony-forming unit–granulocyte/erythrocyte/macrophage/ megakaryocyte.

Tissues ventral to the AGM have an enhancing effect on HSC emergence, while tissues on the dorsal side decrease HSC production ^52, 53^. To study whether there were intrinsic differences in ventrally and dorsally distributed *Mesp1*^der^ PSCs and their capacity to induce EHT, we dissected E11.5 AGM from Mesp1-DsRed^+^ embryos and separated the dorsal and ventral halves ^54^ prior to FACS purification of *Mesp1*^der^ PSCs (Fig. 4B (i)). The CFU-F and replating capacity of the ventral fraction of *Mesp1*^der^ PSCs was comparable to the whole AGM, whereas the dorsal fraction failed to form large colonies and had limited replating capacity (Fig. 4B (ii)-(iii)). Furthermore, whereas the dorsal fraction of *Mesp1*^der^ PSCs failed to induce CFU-Cs from either E11.5 or adult heart endothelium, those from the ventral fraction induced robust numbers including LT-HSCs (Fig. 4B(iv)-(v)). By contrast, co-aggregation of E11.5 or E13.5 *Wnt1*^der^ PSCs with E11.5 (hemogenic) or E13.5 (non-hemogenic) aortic endothelium generated CFU-Cs at very low numbers or not at all, and without any long-term reconstitution when transplanted (Extended Data Fig. 5D (i)-(iii)). CFU-C generation was also comparable across different lots of FCS (Extended Data Fig. 5E).

Taken together, these data show that non-hemogenic embryonic and adult endothelium can be transformed into LT-HSC-producing hemogenic endothelium by co-aggregation with E10.5 or E11.5 *Mesp1*^der^ PSCs, but not with E11.5 or E13.5 *Wnt1*^der^ PSCs.

### *Mesp1*^der^ PSCs induce transcriptomic changes associated with hematopoiesis in non-hemogenic E13.5 endothelium

The cell signalling pathways that coordinate AGM hematopoiesis are not entirely clear, although the roles of WNT ^20^, NOTCH ^19, 55^, and BMP ^21, 22^ signalling in regulating critical transcription factors have been described. Additionally, HSC production is known to be modulated by activation of nitric oxide (NO) synthesis mediated by shear stress and blood flow ^56, 57^, components of prostaglandin E2 ^58^, and inflammatory signals ^59^. Catecholamine signalling in sub-aortic mesenchymal cells has also been shown to impact HSC emergence during embryonic development ^60^.

To explore transcriptional changes in non-hemogenic endothelial cells during their transition to hemogenic endothelium, and to identify features in E11.5 *Mesp1*^der^ PSCs that distinguish them from E11.5 or E13.5 *Wnt1*^der^ PSCs, we performed RNA-sequencing on freshly isolated endothelial cells and *Mesp1*^der^ PSCs, as well as on endothelial cells extracted from co-aggregate cultures at 96 hours (with *Mesp1*^der^ PSCs or without *Mesp1*^der^ PSCs controls). Principal component analysis (PCA) of the transcriptomes of E11.5 and E13.5 endothelial cells extracted from *Mesp1*^der^ PSCs co-aggregates were more closely aligned with each other and with freshly isolated E11.5 hemogenic endothelium than with freshly isolated E13.5 non-hemogenic endothelium or control endothelial cells from 96-hour co-aggregates (Fig. 5A). The PSC fractions were more closely aligned with each other than with endothelial cells; but consistent with their distinct germline derivations and temporal extractions, they were distributed across discrete sectors in transcriptomic space. Consistent with their acquisition of hemogenic properties, genes annotated by Ingenuity Pathway Analysis as associated with development of hematological systems were more differentially expressed in E13.5 endothelial cells extracted from *Mesp1*^der^ PSCs co-aggregates, compared with freshly isolated non-hemogenic E13.5 AGM endothelial cells (Fig. 5B).

**Fig 5.**
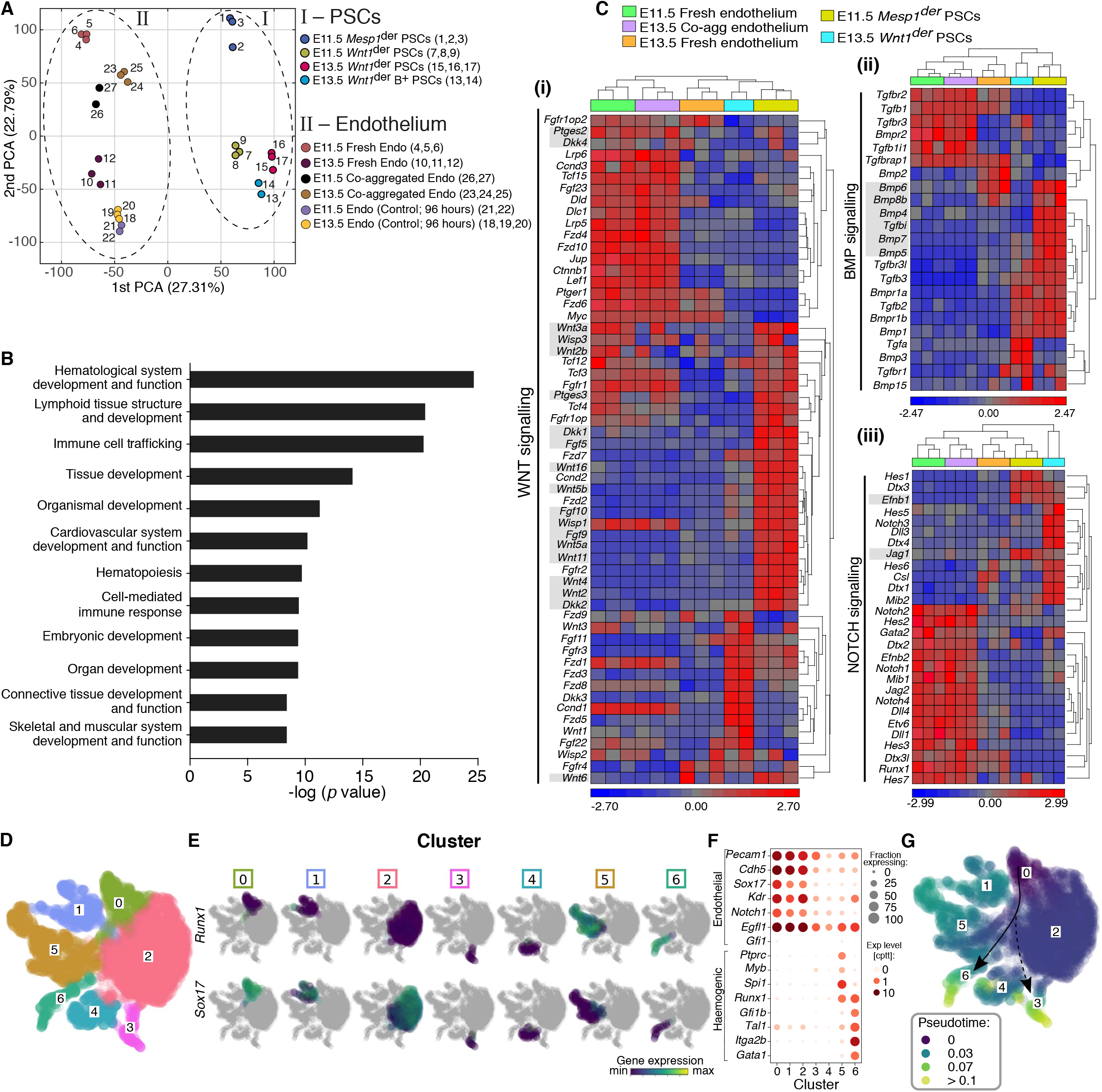
*Mesp1*^der^ PSCs induce transcriptomic changes associated with hematopoiesis in non-hemogenic endothelium. (A) (i) A two-dimensional (2D) principal component analysis (PCA) plot generated from transcriptomes. First, fresh PSCs at E11.5 (*Mesp1*-eYFP^+^ PDGFRA^+^ (*Mesp1*^der^ PSCs)), at E11.5 and E13.5 (*Wnt1*-eYFP^+^ PDGFRA^+^ (*Wnt1*^der^ PSCs)), and at E13.5 (*Wnt1*-eYFP^+^ PDGFRB^+^ (*Wnt1*^der^ B^+^ PSCs)); and second, endothelial cells: fresh E11.5 and E13.5, then E11.5, and E13.5 endothelial cells taken from co-aggregates with E11.5 *Mesp1*-eYFP^+^ PSCs (96 hours), E11.5, and E13.5 control endothelial cells without PSCs (96 hours). (B) Molecular functions annotated by Ingenuity Pathway Analysis of differentially expressed genes: first, in E13.5 endothelial cells co-aggregated with E11.5 *Mesp1*-eYFP^+^ PSCs; and second, in E13.5 control fresh endothelium. (C) Hierarchical clustering of gene expression profiles of hematopoietic mediators associated with (i) WNT signaling, (ii) BMP signaling, and (iii) NOTCH signaling in E11.5 *Mesp1*^der^ PSCs, E13.5 *Wnt1*^der^ PSCs, and E13.5 endothelial cells following co-aggregation with *Mesp1*^der^ PSCs and control E11.5 and E13.5 fresh endothelium. Ligands that were expressed in *Mesp1*^der^ PSCs but not Wnt1der PSCs are highlighted in grey. (D) UMAP representation of single cell transcriptomes from adult cardiac endothelial cells (UBC-GFP mice; GFP^+^, CD31^+^, VE-Cad^+^, PDGFRA^−^) following 0-, 24-, 48-, 72-, 96-hours, co-aggregated with *Mesp1*^der^ PSCs. Endothelial cells from each timepoint were pooled and data from three independent pools are shown together. (E) Distribution of *Runx1* and *Sox17* expression intensities in each cell cluster in (D). (F) Bubble plot representing the distribution of expression levels of various endothelial and hematopoietic genes in each cluster in (D). The size of each bubble represents the fraction of cells in each cluster expressing that gene. (G) UMAP embedding developmental progression of cells in (D) using pseudotime with cluster #0 as the starting reference. Arrows are guides to the eye indicating the expected transition from endothelial to hemogenic cells (solid) and, separately, to a branching transcriptomic profile (cluster 3).

Various components of the WNT ^20^, BMP ^21, 22^, and NOTCH ^55, 61^ signalling pathways have previously been implicated in EHT. We evaluated the expression levels of these components in E11.5 and E13.5 endothelial cells (both freshly isolated and those extracted from *Mesp1*^der^ PSCs co-aggregates in the case of E13.5), E11.5 *Mesp1*^der^ PSCs, and E13.5 *Wnt1*^der^ PSCs (Fig. 5C (i)-(iii)). E11.5 *Mesp1*^der^ PSCs expressed higher levels of several secreted proteins that have previously been associated with EHT, and *Wnt1*^der^ PSCs expressed lower levels. Interestingly, there were higher expression levels in many of the corresponding receptors, signal transducers, and target genes of these ligands in E13.5 endothelial cells extracted from co-aggregate cultures than in freshly isolated non-hemogenic E13.5 endothelial cells, such that they clustered with E11.5 fresh endothelium (Fig. 5C (i)-(iii)). These data suggested a role for WNT, BMP, NOTCH signalling in the induction of EHT in endothelial/*Mesp1*^der^ PSC co-aggregates. To test this, we performed E10.5 and E11.5 *Mesp1*^der^ PSC co-aggregates with adult heart and aortic endothelial cells in the presence of WNT/BMP/NOTCH inhibitors followed by CFU-C and transplantation assays (Extended Data Fig. 6A). With increasing concentrations of WNT inhibitors (WIF-1 or Draxin; Extended Data Fig. 6B (i)), Notch inhibitors (LY450139 or MK0752; Extended Data Fig. 6B (ii)) or a BMP inhibitor (USAG1; Extended Data Fig. 6B (iii)), there was dose dependent inhibition of CFU-Cs and corresponding reduction of donor chimerism in hosts at 6-months following transplantation of aggregates (Extended Data Fig. 6B). Combined inhibition of all three signaling pathways (WIF-1, MK0752 and USAG1) ablated CFU-C and LT-HSC activity (Extended Data Fig. 6C (i)-(ii)).

To capture transcriptional changes in single endothelial cells as they gained hemogenic potential when co-aggregated with AGM stroma, we FAC-sorted adult cardiac endothelial cells from UBC-GFP mice (GFP^+^, CD31^+^, VE-Cad^+^, CD41^−^, CD45^−^, PDGFRA^−,^ PDGFRB^−^) and performed co-aggregate cultures with *Mesp1*^der^ PSCs, FAC-sorted from E11.5 *Mesp1*- DsRed^+^ AGMs (Ds-Red+/PDGFRA+, PDGFRB-, CD31-, VE-Cad-, CD41-, CD45-). Endothelial cells were FAC-sorted (GFP^+^, CD31^+^, VE-Cad^+^, PDGFRA^−^) from 24-, 48, 72, 96-hr co-aggregate cultures, pooled with freshly sorted adult cardiac endothelium and single cell next generation sequencing libraries prepared using a 10X Genomics’ Chromium platform. UMAP representation of endothelial cell transcriptomes from triplicate pools and Leiden clustering detected ^62^ several cell populations (Fig. 5D). SOX17 is expressed in endothelial cells lining arteries but not veins and is a key regulator of hemogenic endothelium ^63^. RUNX1 represses the pre-existing arterial program in hemogenic endothelium and is required for hematopoietic cell transition from hemogenic endothelium ^27, 64^. These transcription factors show a reciprocal expression pattern with most cells in clusters #5 and #6 showing robust *Runx1* expression, and those in clusters #0, #1 and #2, showing high *Sox17* expression (Fig. 5E). Bearing in mind that the endothelial cells were FAC-sorted for sequencing based on PECAM (CD31) and VE-Cad (CDH5) surface protein expression, it is noteworthy that as cells gain *Runx1* expression there was concomitant gain of hematopoietic gene expression (e.g., *Gfi1b*, *Tal1*, *Myb*, *Spi1*, *Gata1*, *Ptprc* (CD45), *Itga2b* (CD41)) and loss of endothelial gene expression (e.g., *Pecam1*, *Cdh5*, *Sox17*, *Kdr*, *Notch1*, *Egfl7*). The clusters with *Runx1* expressing cells appear to have distinct identities with cluster #5 cells showing high levels of *Ptprc* (CD45), *Spi1, Myb*, and those in cluster #6 showing high levels of *Itga2b* (CD41) accompanied by *Gata1, Tal1, Gfi1b* suggestive of distinct hematopoietic lineage differentiation potential prior to complete downregulation of the endothelial signature (Fig. 5F). This is in keeping with reports from single-cell transcriptional analyses of human pluripotent stem cell-derived CD34+ cells ^65^ and pre-HSCs in E11 AGM ^66^. Pseudotime ^67^ (https://arxiv.org/abs/1802.03426) analysis starting from cluster #0 (i.e., strongest endothelial identity; Fig. 5F) showed distinct progression towards cluster #6 (hemogenic) with alternate trajectories towards clusters #3 and #4 (non-hematopoietic) (Fig. 5G).

### PDGFRA-mediated cell signalling is important for LT-HSC generation

The role of *Mesp1*^der^ PSCs in the induction of EHT in co-aggregate cultures prompted us to explore a role for PDGF signalling in mediating endothelial–stromal cell crosstalk in the AGM. The PDGF signalling system consists of four ligands: PDGF-A, -B, -C, and -D ^68^. All four ligands (which are inactive in their monomeric form) assemble intra-cellularly to form disulfide-linked homodimers. PDGF-A and PDGF-B also form heterodimers (PDGF-AB); but because the endogenous expression patterns of PDGF-A and PDGF-B do not frequently overlap, PDGF-AB heterodimers may be infrequent *in vivo*. PDGF-AA binds αα-receptor homodimers ^68^. Expression levels of PDGF-A were highest in E11.5 fresh endothelium, and in E13.5 endothelium harvested from *Mesp1*^der^ PSCs co-aggregates (Fig. 6A (i)). Expression levels of PDGFRA (the receptor for PDGF-A) was highest in E11.5 *Mesp1*^der^ PSCs (Fig. 6A (i)). Confocal images of the E11.5 AGM in *Pdgfra*-nGFP transgenic mice showed abundant and dispersed PDGF-A protein in the endothelium and mesenchyme (Fig. 6A (ii)), but at E13.5, PDGF-A protein levels were low and restricted to the endothelium (Fig. 6A (iii)). Significantly, E11.5 endothelial cells do not express PDGFRA, but sub-aortic stromal cells do (Fig. 6A (i)). To evaluate whether PDGFRA-mediated signalling was required for EHT mediated by *Mesp1*^der^ PSCs at E11.5, we co-aggregated E11.5 *Mesp1*^der^ PSCs with E11.5 aortic endothelium in the presence of a specific PDGFRA inhibitor (APA5) (Fig. 6B (i)). There was dose-dependent reduction in CFU-Cs (Fig. 6B (ii)) and LT-HSCs (Fig. 6B (iii)). A PDGFRB inhibitor (APB5) had no effect on CFU-Cs (Fig. 6B (ii)).

**Fig 6.**
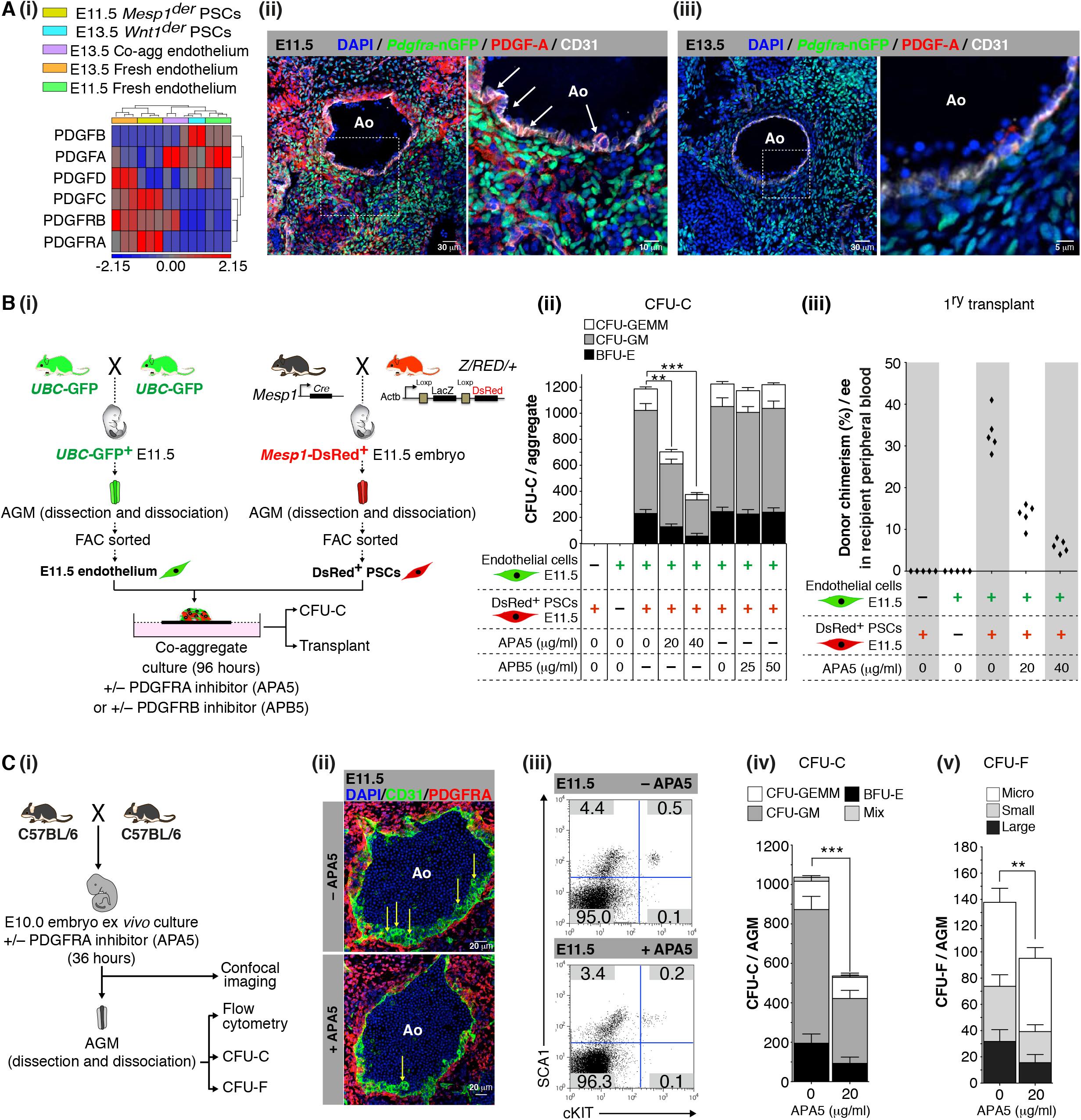
PDGFRA-mediated cell signalling is important for LT-HSC generation. (A) (i) Hierarchical clustering of expression profiles of components of PDGFR signalling in E11.5 *Mesp1*^der^ PSCs, E13.5 *Wnt1*^der^ PSCs, E13.5 endothelial cells following co-aggregation with *Mesp1*^der^ PSCs and control fresh E11.5 and E13.5 endothelium. (ii) Confocal images of an E11.5 *Pdgfra*-nGFP AGM showing high PDGF-A protein in the aortic endothelium and stroma (n=5). (iii) Confocal images of a E13.5 *Pdgfra*-nGFP AGM showing low PDGF-A protein in the aortic endothelium and stroma (n=5). (B) (i) A schematic outlining the process for harvesting cell types used in co-aggregate cultures. Co-aggregate cultures of GFP^+^ endothelial cells (25,000 cells) (*UBC*-GFP^+^/PDGFRA^−^ /PDGFRB^−^/CD31^+^/VECad^+^/CD41^−^/CD45^−^) with DsRed^+^ *Mesp1*^der^ PSCs (150,000 cells) (*Mesp1*-DsRed^+^/PDGFRA^+^/PDGFRB^−^/CD31^−^/VECad^−^/CD41^−^/CD45^−^) in the absence or presence of increasing concentrations of the PDGFRA-specific inhibitor APA5 or PDGFRB-specific inhibitor APB5. (ii) CFU-C analysis of co-aggregate cultures of *Mesp1*^der^ PSCs with E11.5 aortic endothelium at 96 hours. (iii) Percentage of GFP^+^ cells in irradiated recipients (one co-aggregate for each adult irradiated recipient) at six months following transplantation with co-aggregate cultures. (C) (i) A schematic outlining an experiment in which E10.0 embryos were cultured *ex vivo* in the absence or presence of the PDGFRA-specific inhibitor, APA5. (ii) Confocal images of the AGM at E11.5 in the absence (top) or presence (bottom) of APA5 showing no gross architectural changes. (iii) Flow cytometry analysis of cultured E11.5 embryo AGM in the absence (top) or presence (bottom) of APA5, showing reduced SCA1^+^/cKIT^+^ cells in the presence of APA5. (iv) CFU-C analysis of *ex vivo* cultured E11.5 embryo AGM in the absence or presence of APA5. (v) CFU-F analysis of *ex vivo* cultured E11.5 embryo AGM in the absence or presence of APA5. Ao: aortic lumen; CFU-C: colony-forming unit–culture; BFU-E: burst-forming unit–erythroid; CFU-GM colony-forming unit–granulocyte/macrophage; CFU-GEMM: colony-forming unit– granulocyte/erythrocyte/macrophage/megakaryocyte; PDGF-A: platelet-derived growth factor A. CFU-F: colony-forming unit–fibroblast; Micro colonies (<2 mm, 2–24 cells), small colonies (2–4 mm, >25 cells) and large colonies (>4 mm, >100 cells). ** p<0.01 and *** p<0.001; unpaired two-tailed t-test. Data represent mean ± s.e.m.

To investigate the impact of PDGF-A/PDGFRA signalling on EHT *in vivo*, we harvested E10.0 embryos and cultured them *ex vivo* ^69, 70^ in the presence or absence of the PDGFRA inhibitor-APA5 (Fig. 6C (i)). After 36 hours we evaluated the AGM for structural changes by confocal microscopy, and the impact on hematopoiesis and PSCs by flow cytometry, CFU-C, and CFU-F assays. Confocal immunofluorescence imaging showed that inhibition of PDGFRA signalling at E10 did not severely impact on the architecture of the dorsal aorta, but blood clusters along its ventrolateral surface appeared reduced (Fig. 6C (ii)). Evaluating this further by flow cytometry showed that SCA1^+^/CKIT^+^ and SCA1^+^/CKIT^−^ cells were reduced by 60% and 22% respectively (Fig. 6C (iii)). Both CFU-Cs (Fig. 6C (iv)) and CFU-Fs (Fig. 6C (v)) were significantly reduced in number. Conversely, PDGF-AA supplementation of media lacking the cytokine cocktail (IL3, SCF, Flt3L) that is used in co-aggregate cultures, rescued CFU-C and LT-HSC production from endothelial cells when co-aggregated with *Mesp1*^der^ PSCs but had no impact on *Wnt1*der PSCs (Extended Data Fig. 7A (i)-(iii)).

As summarised in Fig. 7, the aortic endothelium is derived in part from E9.5 *Mesp1*^der^ PSCs, which at E11.5 are distributed in the aortic sub-endothelium and express WNT/NOTCH/BMP hemogenic ligands. The E11.5 endothelium has complementary receptors for these ligands and produce soluble PDGF-AA. Interrupting PDGF-AA/PDGFRA signalling impedes *Mesp1*^der^ PSCs function and EHT. Coincident with the cessation of EHT at E13.5, sub-endothelial PDGFRA^+^ *Mesp1*^der^ PSCs are replaced by *Wnt1*^der^ PSCs. *Wnt1*^der^ PSCs do not express hemogenic ligands. The non-hemogenic E13.5 endothelium also does not express complementary receptors for these ligands or soluble PDGF-AA. However, as shown here, EHT can be re-established in non-hemogenic endothelium by co-aggregating these cells with E10.5 or E11.5 *Mesp1*^der^ PSCs to generate LT-HSCs.

**Fig 7.**
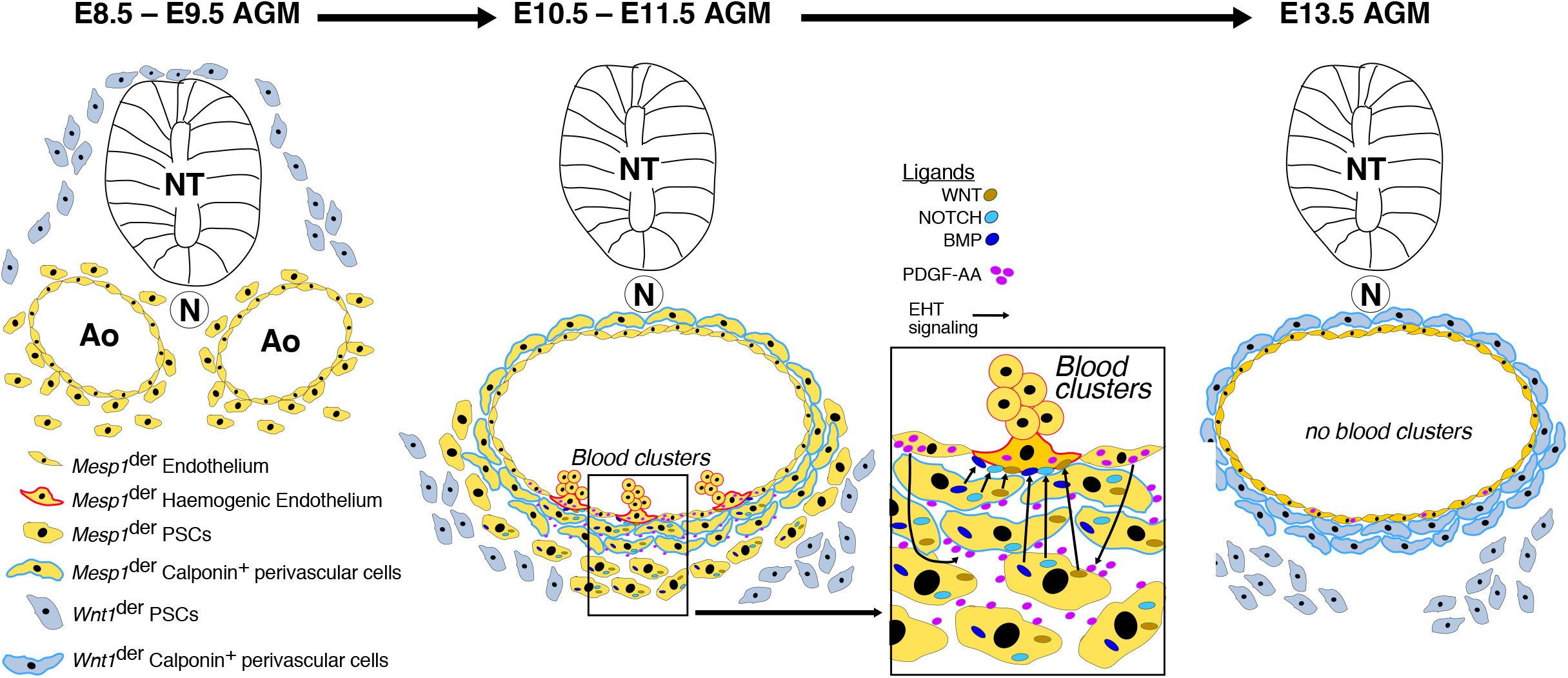
Proposed model summarising the roles of *Mesp1*^der^ PSCs in HSC development in the AGM. A model that incorporates the changing landscape of *Mesp1*- and *Wnt1*-derived PSCs in the AGM stroma, and their role in generating endothelial and sub-endothelial cells, and LT-HSCs. Pre- E9.5/E9.5 *Mesp1^der^* PSCs contribute to the aortic endothelium. E10.5/E11.5 endothelium secretes PDGF-AA, which acts on PSCs that secrete hemogenic factors to promote EHT. *Mesp1^der^* PSC are replaced by *Wnt1^der^* PSCs at E13.5. This is accompanied by the loss of high PDGF-AA in the AGM and interruption of a PDGFRA mediated signaling axis involving *Mesp1^der^* PSC dependent induction of EHT.

## DISCUSSION

We have identified a transient population of *Mesp1*^der^ PSCs in the mouse AGM that can induce EHT and generate LT-HSCs when co-aggregated with endothelial cells. These multipotent and clonogenic PSCs, which have long-term growth potential *in vitro*, are distributed around the dorsal aorta and contribute to its structures, including the endothelial, pericyte, and smooth muscle layers. Ventrally distributed *Mesp1*^der^ PSCs have high CFU-F and long-term growth potential and cooperate with endothelial cells to induce EHT and LT-HSC generation in the AGM.

PDGFRA^+^ is broadly expressed in primitive para-axial mesoderm, somites, and mesenchyme ^71^ and contributes to hemogenic endothelium and HPCs ^36^. The PDGFRA^+^/Flk1^+^ population has been proposed as the source for hemato-endothelial lineages from the PDGFRA^+^ population. PDGFRA^+^/Flk1^+^ cells persist to E9.5 but at significantly reduced numbers ^36^. Here we show that AGM PDGFRA^+^ stromal cells, especially those that lack pericyte or endothelial commitment, have long-term replating capacity. These cells can act as a reservoir that seed these lineages and continue to contribute to hemogenic endothelium to E9.5.

PDGF-AA/PDGFRA signalling plays a major role in stromal-endothelial cell crosstalk. At E11.5 the aortic endothelium expresses abundant levels of PDGF-AA, which diffuses through and permeates cells in the sub-endothelial and deeper stromal layers. These layers in turn express robust levels of the cognate receptor, PDGFRA. Notably, endothelial cells do not themselves express PDGFRA and at E13.5 they show sparse PDGF-AA ligand expression. It has been proposed that hematopoietic production at this site depends on a limited, non-renewable, pool of hemogenic endothelial cells that are replaced by somite-derived endothelial cells and that this may account for the cessation of AGM hematopoiesis at E13.5 ^72, 73^. Whether replacement of *Mesp1*^der^ PSCs, which promote EHT, by *Wnt1*^der^ PSCs that do not, also contributes to cessation of AGM hematopoiesis is an intriguing possibility.

MSCs populate embryonic tissues in waves, with Sox1^+^ neuroepithelium- rather than mesoderm-derived MSCs being the earliest to populate the trunk ^35^. But these cells were reported to be transient and are replaced by MSCs from an unknown source later during development ^35^. Here, we examined stromal cell populations at earlier embryonic time-points and focused on the AGM, which has a well-defined role in HSC generation. We have shown that at E11.5, *Mesp1*^der^ PSCs contact the aortic endothelium directly, with *Wnt1*^der^ PSCs located 5 or 6 cell layers deeper in the stroma. At E13.5, *Mesp1*^der^ PSCs are no longer evident, and *Wnt1*^der^ PSCs are now in direct contact with endothelium at the ventral surface of the dorsal aorta. What accounts for the disappearance of *Mesp1*^der^ PSCs from the AGM at E13.5 is not known. It is possible that these cells stopped proliferating and went through terminal differentiation as a new wave of PSCs infiltrated the AGM. Alternatively, these cells could have undergone apoptosis, or migrated elsewhere in the embryo. There are precedents to distinct sub-populations of MSCs contributing to specific tissues during development ^74^. Proliferative mesoderm-derived NESTIN^−^ MSCs participate in fetal skeletogenesis and lose MSC activity soon after birth. In contrast, quiescent neural crest-derived NESTIN^+^ cells do not generate fetal chondrocytes but preserve MSC activity and later differentiate into HSC niche-forming MSCs, helping to establish the bone marrow HSC niche by secreting CXCL12 ^74^.

The transcriptomes of *Mesp1*^der^ PSCs and *Wnt1*^der^ PSCs are distinct, and expression of high levels of NOTCH, BMP, and WNT ligands in the former are consistent with their role in hematopoiesis. It is indeed likely that *Mesp1*^der^ PSCs are themselves a heterogeneous population of cells evident by intrinsic functional differences between cells located in the dorsal or ventral halves of the AGM (Fig. 4B). Single-cell transcriptional analyses of *Mesp1*^der^ PSCs may help resolve this heterogeneity. It is a tool that could also be used to identify *Mesp1*^der^ PSC sub-populations in E9.5 (and earlier), which might be committing to endothelial and/or perivascular cell fates and others that remain in a mesenchymal state to promote EHT at E10.5/E11.5. Particularly striking is the induction of receptors for BMP, WNT, NOTCH ligands in non-hemogenic E13.5 endothelial cells at levels comparable to those of E11.5 endothelium in co-aggregate cultures with *Mesp1*^der^ PSCs; and this was not observed in *Wnt1*^der^ PSCs. The factors that drive expression of these receptors are not known, but our data show a distinct role for PDGF-AA/PDGFRA signalling in the induction, by *Mesp1*^der^ PSCs, of LT-HSC production in endothelial cells. Identifying equivalent PSCs in the human AGM or factors that could replace the use of embryonic PSCs altogether, are key steps toward generating autologous HSCs from human endothelial cells for disease modelling and therapy. Compared to methods that rely on introducing transcription factors to mediate cell-fate conversion ^4, 5^, these data point to a more convenient solution for the generation of hematopoietic stem cells.

The single-cell transcriptomes of endothelial cells when co-aggregated with *Mesp1*^der^ PSCs revealed several cell clusters with high endothelial gene expression and others that appear to lose their endothelial identity as they activated hematopoietic gene expression. There were others that were low in both endothelial and hematopoietic gene expression. Although the precise developmental trajectory of transitioning cells is currently not known, pseudotime analysis revealed a distinct trajectory from mature endothelial cells to hemogenic endothelial cells. Interestingly, this analysis also revealed an alternate trajectory to non-hemogenic endothelial cell clusters. Whether these cells are by-products or necessary for the generation of hemogenic endothelial cells is currently not known. However, these data can be used to guide the prospective isolation of relevant cell populations to experimentally validate these trajectories. This would enable comparison with single cell transcriptomic data from primary pre-hematopoietic cells in the AGM ^44, 75^ to establish whether the steps followed in adult endothelial cell reprogramming by *Mesp1*^der^ PSCs are comparable with those in the AGM.

Taken together, our findings identify a population of PSCs in the mouse AGM that induce LT-HSC production in endothelial cells. These PSCs constitute a trackable system for interrogating the molecular processes that drive EHT, and a tool to potentially generate engraftable LT-HSCs for the treatment of hematological disorders.

## ONLINE CONTENT

Methods, along with additional Extended Data display items and Source Data, are available in the online version of the paper; references unique to those sections appear only in the online paper.

## ACKNOWLEDGMENTS

Professor W. D. Richardson, University College London and Dr K. M. Young, University of Tasmania for providing *PdgfraCre*^ERT2^ reporter mice. Professor A. Waisman, Johannes Gutenberg University Mainz for providing iDTR mice. Professor R. P. Harvey Victor Chang Cardiac Research Institute, Sydney for transferring *Pdgfra-nGFP*, *Wnt1Cre*, *Mesp1Cre* mice. UNSW Sydney Biological Resource Centre staff for maintaining mouse lines, and the Mark Wainwright Analytical Centre for assistance with flow cytometry, confocal microscopy and image processing. The authors gratefully acknowledge discussions with Dr Nancy Speck (University of Pennsylvania, US), Dr Alexander Medvinsky and Dr Elaine Dzierzak (University of Edinburgh, UK). This work was funded by the National Health and Medical Research Council of Australia (J.E.P., 510100, 568668, 630497, 1102589; and V.C., 1061593), Australian Research Council (J.E.P., DP0984701), Faulty of Medicine, UNSW Grant (V.C), and a Mark Wainwright Analytical Centre, UNSW Grant (V.C.). BG was supported by Wellcome (206328/Z/17/Z), MRC (MR/S036113/1), Blood Cancer UK (#18002) and core funding by Wellcome to the Cambridge Stem Cell Institute.

## SUPPLEMENTARY INFORMATION

See online content

## AUTHOR CONTRIBUTIONS

V.C. and J.E.P. designed the study. V.C. performed most of the experiments and analysed the data for the manuscript. P.R., Y.C.K., K.K., Q.Q., R.A.O, AU, B.L., C.B. and C.P. performed experiments. F.Z., Y.H., D.C., D.R.C., and D.B. performed bioinformatics analysis on the sequencing data and B.G. with data interpretation. S.M., G.E., W.W., and S.T. contributed valuable tools and protocols. V.C., F.Z., B.G. and J.E.P. interpreted results. V.C. and J.E.P. wrote the manuscript. All authors have read and approved the manuscript.

## EXTENDED FIGURES

**Extended Data Fig. 1:**
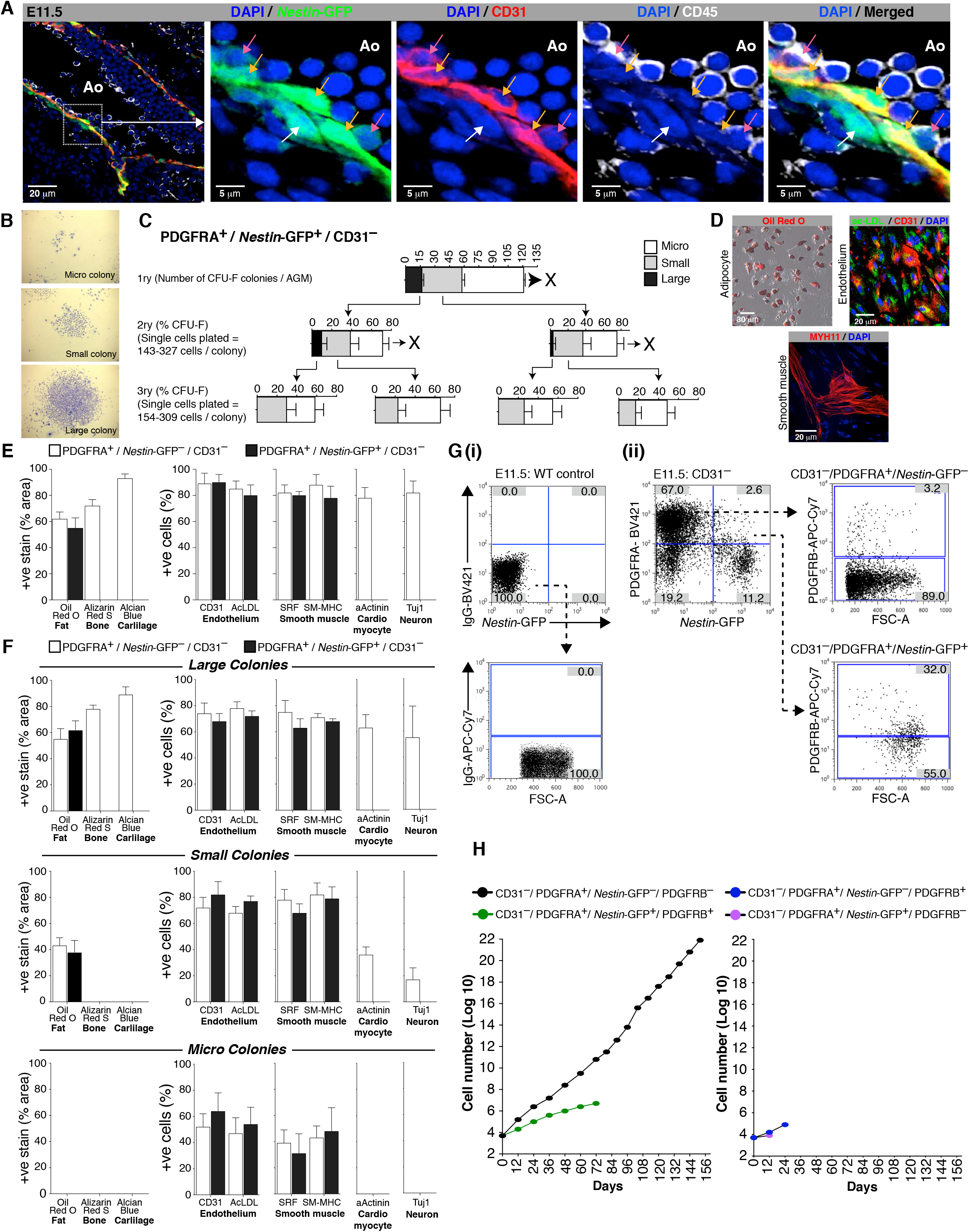
The E11.5 AGM has resident long- and short-term repopulating CFU-Fs. (A) Confocal images of the ventral surface of an E11.5 dorsal aorta showing *Nestin*-GFP (green) expression in CD31^+^ endothelial (orange arrow), CD31^−^ perivascular mesenchymal (white arrow) and budding CD45^+^ blood cells (pink arrow). (B) CFU-F classification based on size and cell number. (C) Single cell clonal analysis of CD31^−^; PDGFRA^+^; *Nestin*-GFP^+^ CFU-Fs. (D) *In vitro* differentiation of CD31^−^; PDGFRA^+^; *Nestin*-GFP^+^ cells (n=3). (E) Bar graphs showing % (percentage) conversion to marker-positive cells following differentiation induction of bulk cultured CD31^−^; PDGFRA^+^; *Nestin*-GFP^−^ and CD31^−^; PDGFRA^+^; *Nestin*- GFP^+^ CFU-F cells. (F) Bar graphs showing % (percentage) conversion to marker-positive cells following differentiation induction of CD31^−^; PDGFRA^+^; *Nestin*-GFP^−^ and CD31^−^; PDGFRA^+^; *Nestin*-GFP^+^ single cell derived large-, small- and micro- CFU-F colonies. (G) (i) Isotype staining controls. (ii) Percentage of E11.5 *Nestin*-GFP^+^ AGM CD31^−^; PDGFRA^+^; *Nestin*-GFP^−^ and CD31^−^; PDGFRA^+^; *Nestin*-GFP^+^ cells that also express PDGFRB. (H) Long-term growth of E11.5 AGM-derived CFU-Fs based on CD31, PDGFRA, *Nestin*-GFP and PDGFRB expression (average values from n=3). Ao: aortic lumen; CFU-F: colony-forming unit–fibroblast; colony sizes: micro colonies (<2 mm, 2–24 cells), small colonies (2–4 mm, >25 cells) and large colonies (>4 mm, >100 cells).

**Extended Data Fig. 2:**
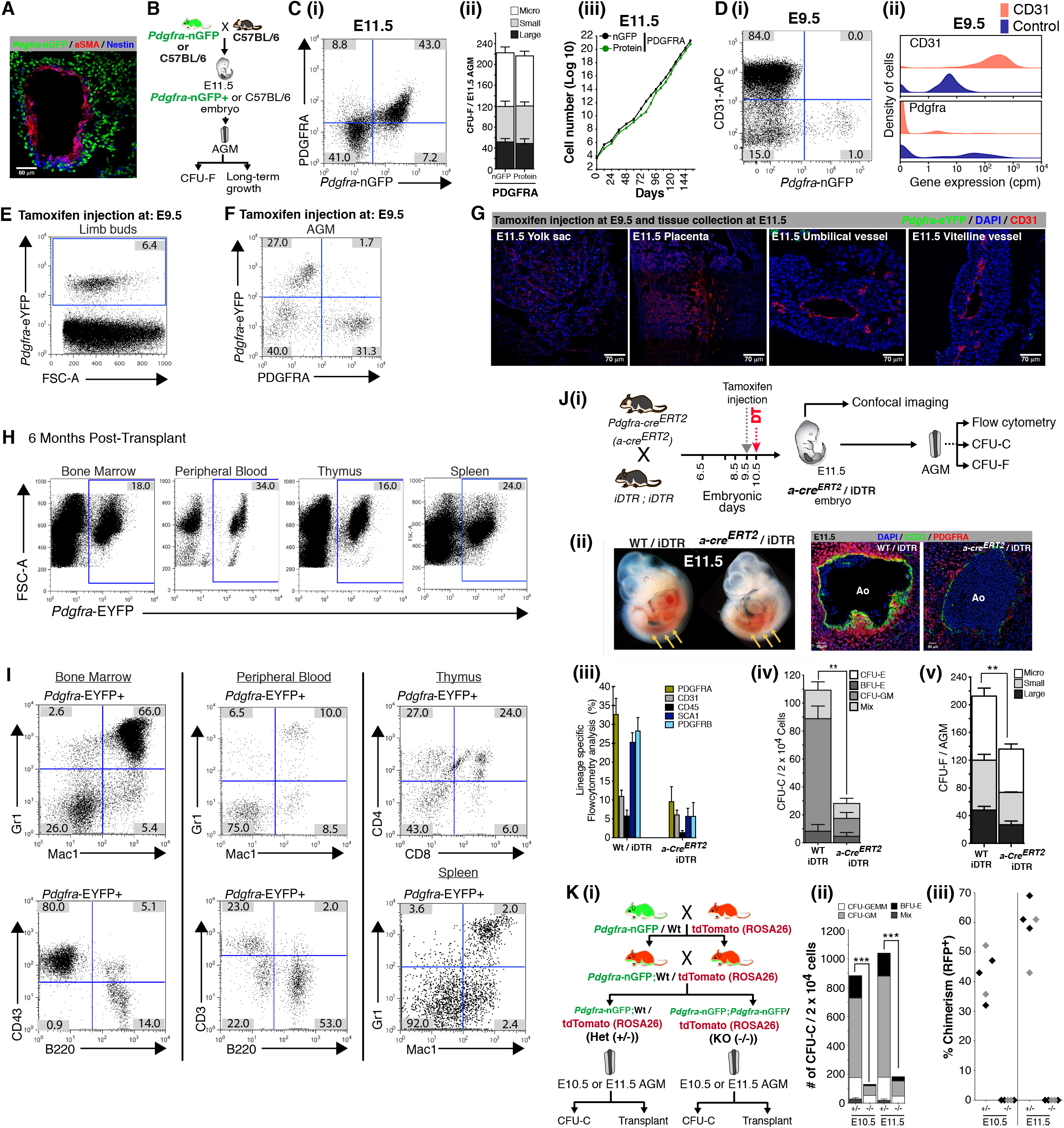
Characterisation of PDGFRA^+^ cell contributions to the developing aorta and LT-HSCs. (A) Spatial distribution of *Pdgfra*-nGFP, NESTIN, and aSMA expressing cells in a *Pdgfra*-nGFP E11.5 AGM. (B) Schematic outline of experiments using E11.5 wild-type or *Pdgfra*-GFP^+^ AGM to evaluate comparability of PDGFRA protein and *Pdgfra*-nGFP positivity. (C) (i) Flow cytometry of E11.5 AGMs harvested from *Pdgfra*-nGFP embryos. (ii) Comparison of CFU-F potential of cells expressing *Pdgfra*-nGFP and cells expressing PDGFRA protein in the E11.5 AGM (n=5). (iii) Long-term growth of *Pdgfra*-nGFP and PDGFRA protein expressing CFU-Fs (averaged values from n=3). (D) (i) Flow cytometry of E9.5 *Pdgfra*-nGFP^+^ AGM for CD31 and GFP expression. (ii) CD31 and Pdgfra expression levels in single CD45^−^, CD31^+^, CD144^+^ endothelial and single CD45^−^, CD31^−^, CD144^−^ control cells sorted from the E9.5 embryo proper excluding head, limb buds, heart and visceral bud (PMID 32203131). (E) Flow cytometry of limb buds harvested from E11.5 *Pdgfra-cre^ERT2^ R26R*-eYFP embryos after a single injection of tamoxifen at E9.5. (F) Flow cytometry of AGMs harvested from E11.5 *Pdgfra-cre^ERT2^ R26R*-eYFP embryos after a single injection of tamoxifen at E9.5. (G) Confocal images of E11.5 *Pdgfra-cre^ERT2^ R26R*-eYFP embryo yolk sac, placenta, umbilical and vitelline vessels after a single injection of tamoxifen at E9.5. (H) Flow cytometry analysis of donor eYFP^+^ cells (from neonatal bone marrow following *cre*-activation at E9.5) in various tissues in primary transplants. (I) (i) Flow cytometry analysis of donor eYFP^+^ cell (from neonatal bone marrow following *cre*-activation at E9.5) contributions to various blood cell fractions in the bone marrow, peripheral blood, thymus, and spleens in recipients 6 months post-transplantation. (J) (i) Schematic showing the experimental strategy used to ablate PDGFRA^+^ cells. (ii) Phenotypic changes in E11.5 wild-type – and PDGFRA^+^ – cell-ablated whole embryos (arrows indicate the AGM region). Confocal images of control (Wt / iDTR) (left) and *Pdgfra*-*cre^ERT2^ /* iDTR (right) E11.5 AGMs following tamoxifen induction at E9.5 and diphtheria toxin treatment at E10.5. (iii) Flow cytometry of control (Wt / iDTR) and *Pdgfra*-*cre^ERT2^* / iDTR E11.5 AGM (following tamoxifen induction at E9.5 and diphtheria toxin treatment at E10.5) to quantify numbers of blood (CD45), endothelial (CD31), pericyte (PDGFRB), and CFU-Fs (PDGFRA). (iv) CFU-C quantification in E11.5 AGM (n=14) after tamoxifen-induced PDGFRA^+^ cell ablation. (v) CFU-Fs in E11.5 AGM (n=12) after tamoxifen-induced PDGFRA^+^ cell ablation. (K) (i) Schematic showing the experimental strategy used to generate *Pdgfra*-nGFP knockout (KO) embryos. (ii) CFU-Cs in *Pdgfra* knockout E10.5 and E11.5 AGM (n=11). (iii) Flow cytometry analysis of donor tdTomato^+^ cells (from *Pdgfra* knockout and wildtype AGM) in peripheral blood of recipient mice at six months post-bone marrow transplantation. CFU-F: colony-forming unit–fibroblast; colony sizes: micro colonies (<2 mm, 2–24 cells), small colonies (2–4 mm, >25 cells) and large colonies (>4 mm, >100 cells). FSC-A: forward scatter area. Unpaired two-tailed t-test. Data represent mean ± s.e.m. Ao: aortic lumen; CFU-C: colony-forming unit–culture; CFU-E: colony-forming unit–erythroid; BFU-E: burst-forming unit–erythroid; CFU-GM colony-forming unit–granulocyte/macrophage; CFU-F: colony-forming unit–fibroblast; colony sizes: micro colonies (<2 mm, 2–24 cells), small colonies (2–4 mm, >25 cells) and large colonies (>4 mm, >100 cells). *Pdgfra* KI/KI (null): –/– and *Pdgfra* KI/+ (heterozygote): +/–.

**Extended Data Fig. 3:**
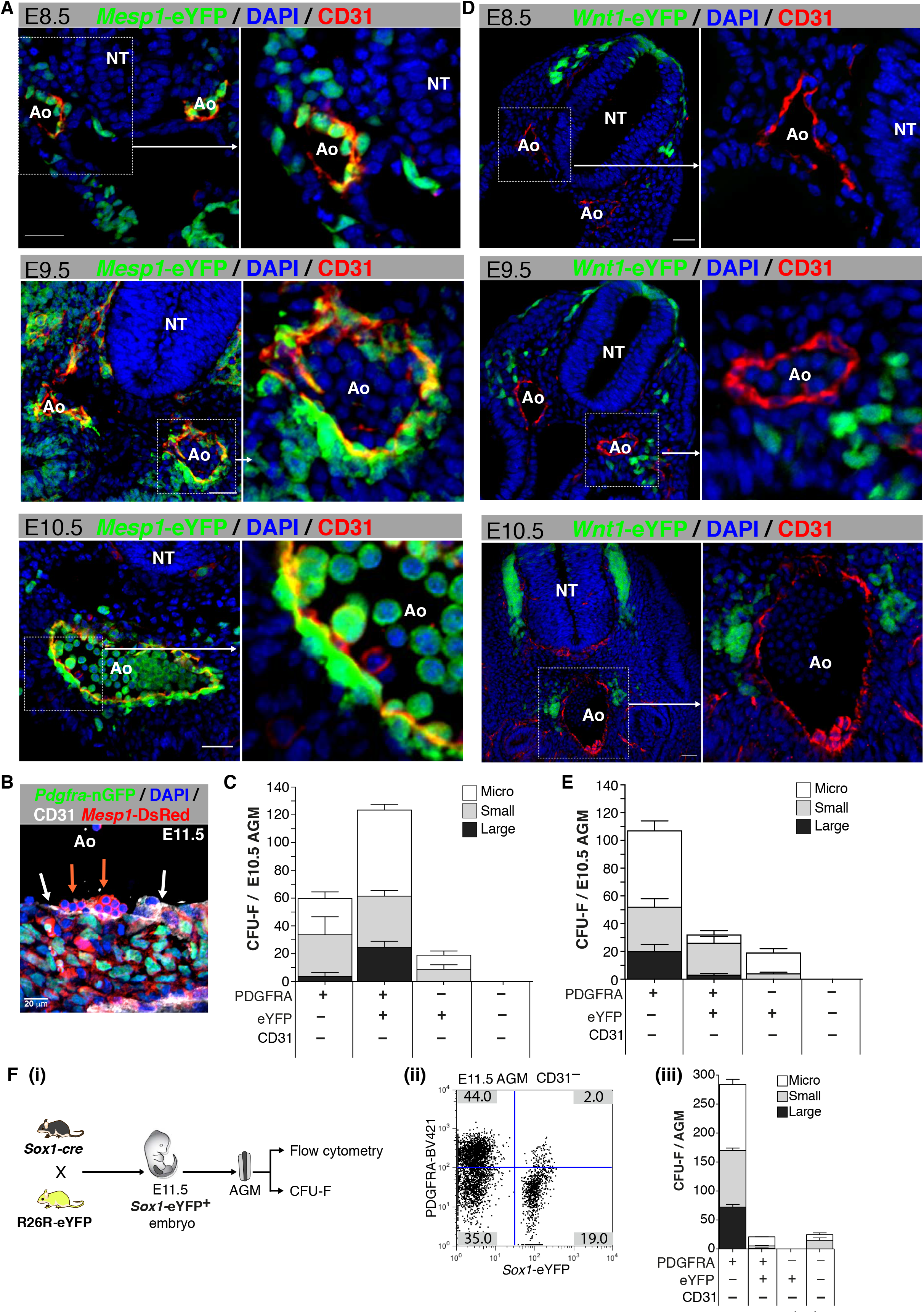
Germ layer contributions to PDGFRA^+^ PSCs in the AGM. (A) (i) Confocal images of E8.5, E9.5 and E10.5 AGMs in *Mesp1*-eYFP embryos. (B) Confocal image of the ventral surface of a E11.5 dorsal aorta in a *Pdgfra*-nGFP*/Mesp1*-DsRed double transgenic embryo (n=4) showing DsRed^+^ blood (orange arrows), endothelium (white arrows), and nGFP^+^/ DsRed^+^ stromal cells. (C) CFU-Fs in cell fractions sorted from *Mesp1*-eYFP^+^ AGMs (n=9) at E10.5. (D) Confocal images of E8.5, E9.5 and E10.5 AGMs in *Wnt1*-eYFP embryos. (E) CFU-Fs in cell fractions sorted from *Wnt1*- eYFP^+^ AGMs (n=12) at E10.5. (F) (i) A schematic outline of the genetic cross used to generate and harvest *Sox1*-eYFP embryos (n=12) at E11.5. (ii) Flow cytometry showing the percentage of *Sox1*-eYFP cells in the E11.5 AGM. (iii) CFU-Fs in *Sox1*-eYFP E11.5 AGM. Ao: aortic lumen; DAPI: 4’,6-diamidino-2-phenylindole dihydrochloride; BV421: Brilliant Violet 421; CFU-F: colony-forming unit–fibroblast; colony sizes: Micro colonies (<2 mm, 2–24 cells), small colonies (2–4 mm, >25 cells) and large colonies (>4 mm, >100 cells). Data represent mean ± s.e.m.

**Extended Data Fig. 4:**
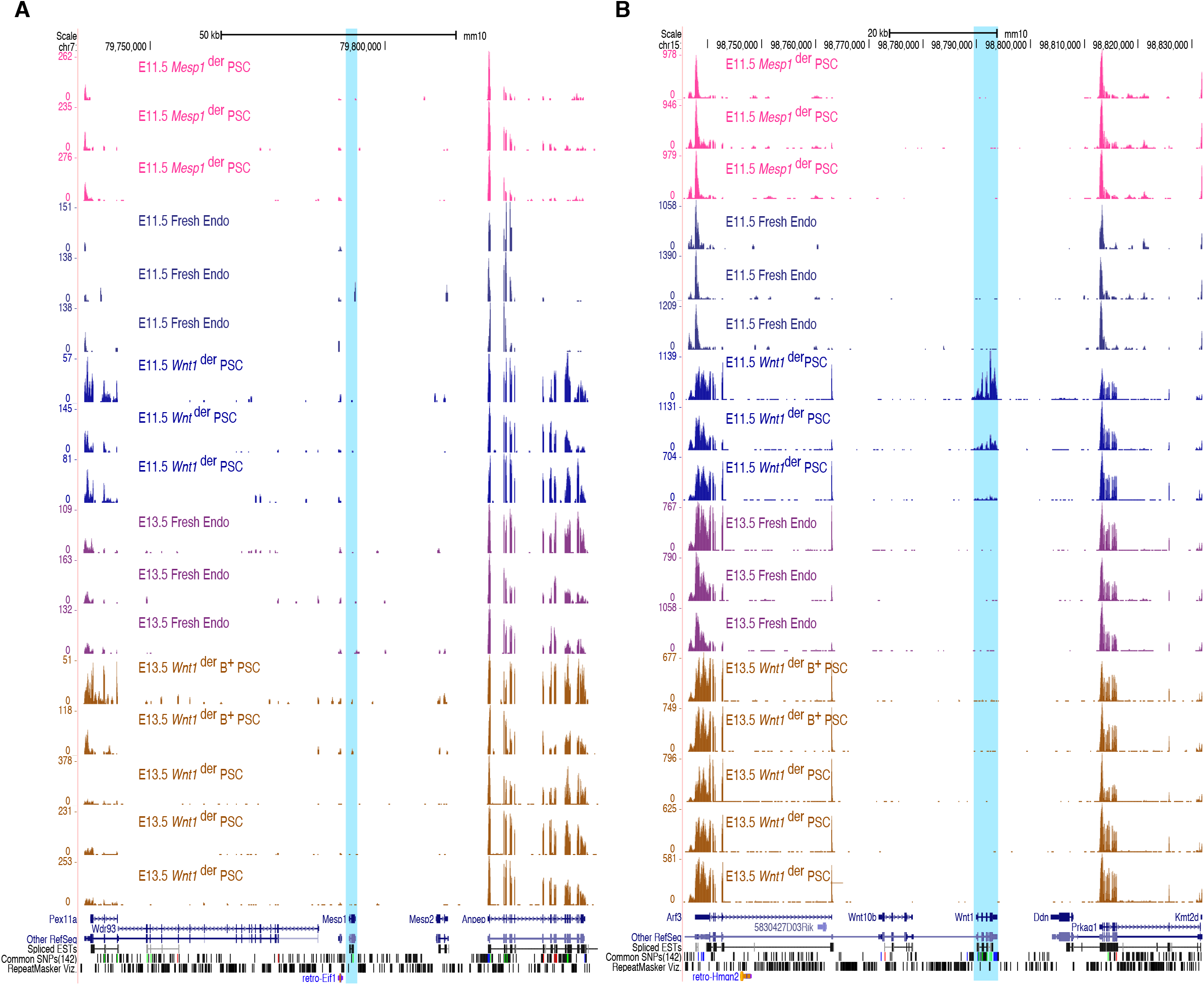
Evaluation of *Mesp1* and *Wnt1* transcripts in E11.5 and E13.5 AGM cell subsets. (A) Evaluation of *Mesp1* expression in AGM cell-subsets. RNA sequencing traces mapped to the *Mesp1* locus and its flanking genes. *Mesp1*^der^ PSCs: *Mesp1*- eYFP^+^/PDGFRA^+^/PDGFRB^−^/CD31^−^/VE-Cad^−^/CD41^−^/CD45^−^, *Wnt1*^der^ PSCs: *Wnt1*-eYFP^+^/ PDGFRA^+^/PDGFRB^−^/CD31^−^/VE-Cad^−^/CD41^−^/CD45^−^, *Wnt1*^der^ B^+^ PSCs: *Wnt1*-eYFP^+^/ PDGFRA^+^/PDGFRB^+^/CD31^−^/VE-Cad^−^/CD41^−^/CD45^−^ and Endo: endothelial cells; PDGFRA^−^ /PDGFRB^−^/CD31^+^/VE-Cad^+^/CD41^−^/CD45^−^. (B) Evaluation of *Wnt1* expression in AGM cell subsets. RNA sequencing traces mapped to the *Wnt1* locus and its flanking genes.

**Extended Data Fig. 5:**
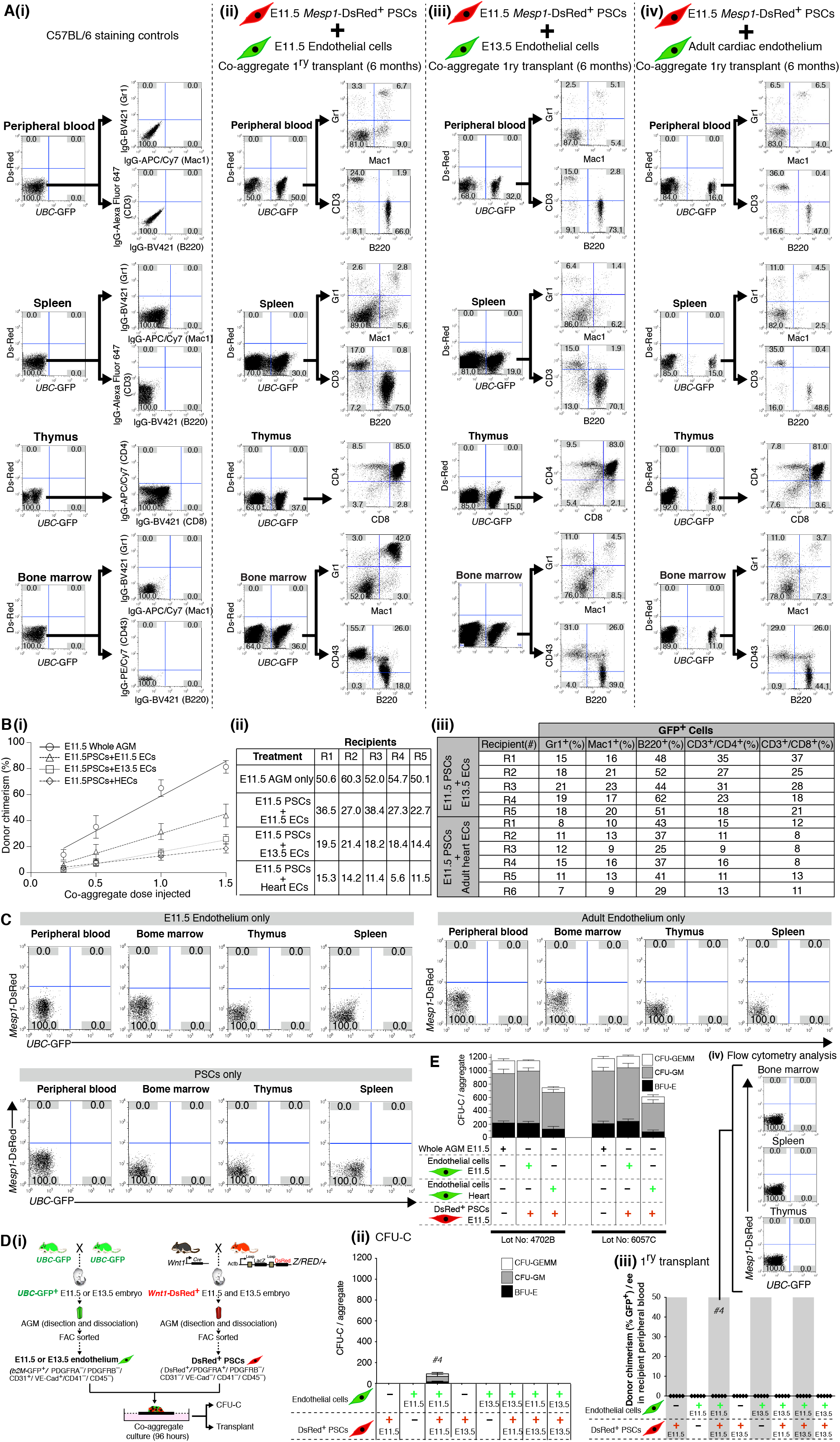
Multilineage reconstitution from embryonic and adult heart endothelial cell derived LT-HSCs. (A) (i) Staining controls using adult C57BL/6 wild-type peripheral blood, spleen, thymus and bone marrow. (ii) Flow cytometry using hematopoietic tissues from recipients six months after transplantation with E11.5 endothelium and E11.5 *Mesp1*^der^ PSC co- aggregates. (iii) Flow cytometry using hematopoietic tissues from recipients six months after transplantation with E13.5 endothelium and E11.5 *Mesp1*^der^ PSC co-aggregates and (iv) Flow cytometry using hematopoietic tissues from recipients six months after transplantation with adult cardiac endothelium and E11.5 *Mesp1*^der^ PSC co-aggregates. (B) (i) Relative donor chimerism and HSC numbers from whole AGM reaggregates and PSC/endothelial co-aggregates (ii) Table summarising HSC numbers calculated from serial dilutions/transplantation. (iii) Table summarising the contribution of donor cells to various blood lineages in recipient mice. (C) Flow cytometry using hematopoietic tissues from recipients six months after transplantation with re-aggregates of E11.5 endothelium or adult cardiac endothelium or E11.5 PSCs. (D) (i) Schematic diagram showing the experimental strategy used to generate cells for co-aggregation of endothelial cells with *Wnt1^der^* PSCs. (ii) CFU-C assays from 96 h co-aggregate cultures (n=5–7 co-aggregates per culture condition). (iii) Percentage of GFP^+^ cells in peripheral blood of recipients (n=5 co-aggregates per culture condition). One co-aggregate per recipient mouse and each point represents an individual recipient. (iv) Flow cytometry analysis of bone marrow, spleen, and thymus for GFP^+^ cells. PSC; PDGFRA+ stromal cells, AGM; aorta gonad mesonephros, EC; endothelial cell, HEC; adult heart endothelial cells. (E) CFU-C analysis of co-aggregate cultures using two lots of FCS. CFU-C: colony-forming unit–culture; BFU-E: burst-forming unit–erythroid; CFU-GM; colony-forming unit– granulocyte/macrophage; CFU-GEMM: colony-forming unit– granulocyte/erythrocyte/macrophage/megakaryocyte. Data represent mean ± s.e.m.

**Extended Data Fig. 6:**
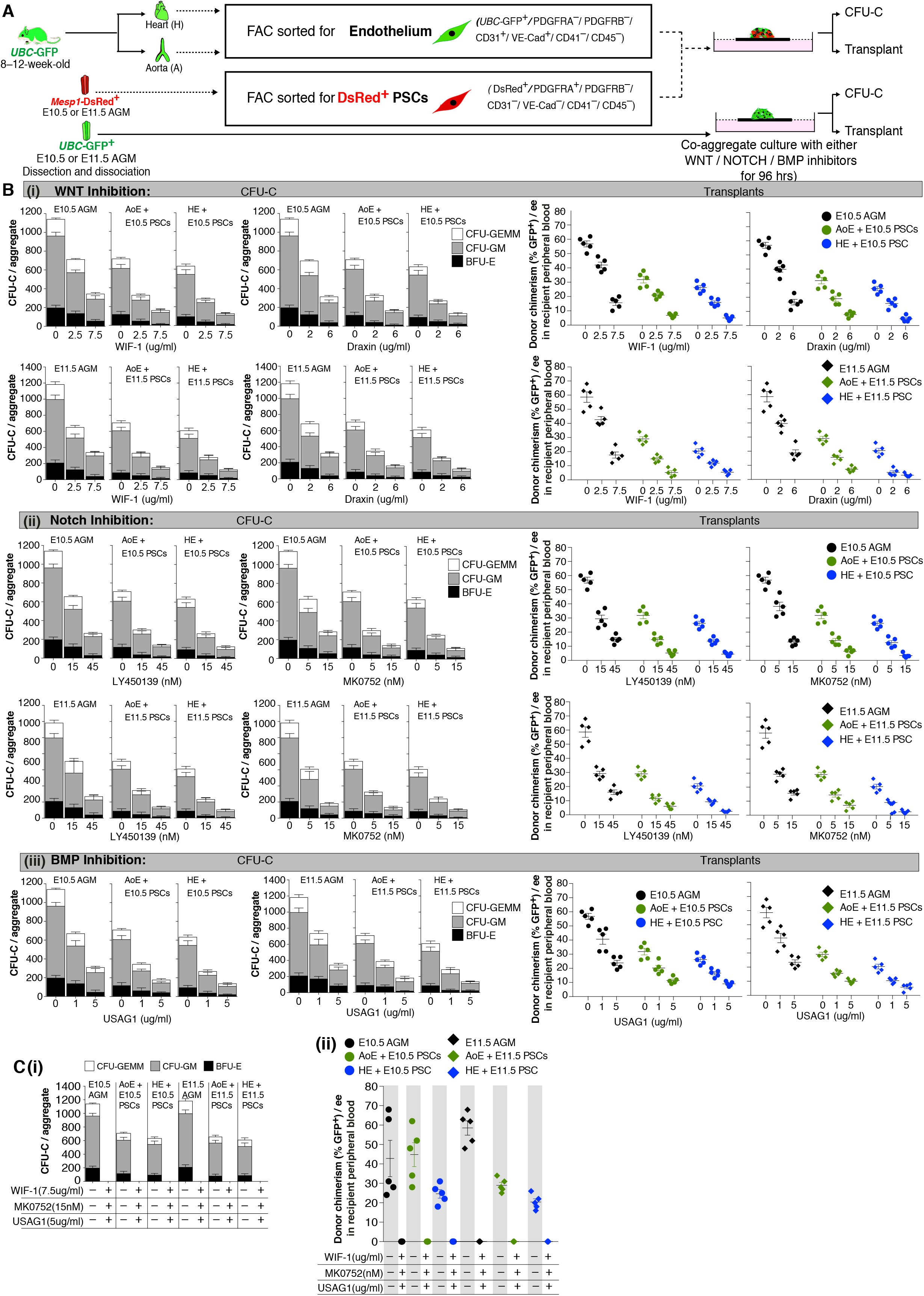
Conversion of adult endothelial cells into HSC producing hemogenic endothelium requires WNT, NOTCH and BMP signalling. (A) Schematic outlining the process for harvesting cell types used in co-aggregate cultures with and without specific WNT, NOTCH and BMP inhibitors. PSCs (150,000 cells) from E10.5 and E11.5 embryos were co-aggregated with single cell suspensions from adult heart endothelium (25,000 cells), adult aortic endothelium (25,000 cells) or re-aggregated whole AGM and cultured for 96 hours. The PSCs were DsRed^+^ (*Mesp1*- DsRed^+^/PDGFRA^+^/PDGFRB^−^/CD31^−^/VECad^−^/ CD41^−^/CD45^−^); the endothelial cells were GFP^+^ (*UBC*-GFP^+^/PDGFRA^−^/PDGFRB^−^/CD31^+^/VECad^+^/CD41^−^/CD45). (B) CFU-C potential (left) and donor chimerism at six months following bone marrow transplantation (right; one aggregate per irradiated adult recipient) of E10.5 or E11.5 AGM re-aggregates and adult aortic endothelium (AoE) or adult heart endothelium (HE) co-aggregated with E10.5 or E11.5 *Mesp1*-DsRed^+^ PSCs in the presence of increasing concentrations of (i) WNT inhibitors, (ii) NOTCH inhibitors and (iii) a BMP inhibitor. (D) (i) CFU-C potential of E10.5 or E11.5 AGM re-aggregates or adult aortic endothelium (AoE) and adult heart endothelium (HE) co-aggregated with E10.5 or E11.5 *Mesp1*-DsRed^+^ PSCs in the presence of all three (WNT, NOTCH and BMP) inhibitors. (ii) donor chimerism at six months following bone marrow transplantation (one aggregate per irradiated adult recipient) of reaggregates and co-aggregates from D (i).

**Extended Data Fig. 7:**
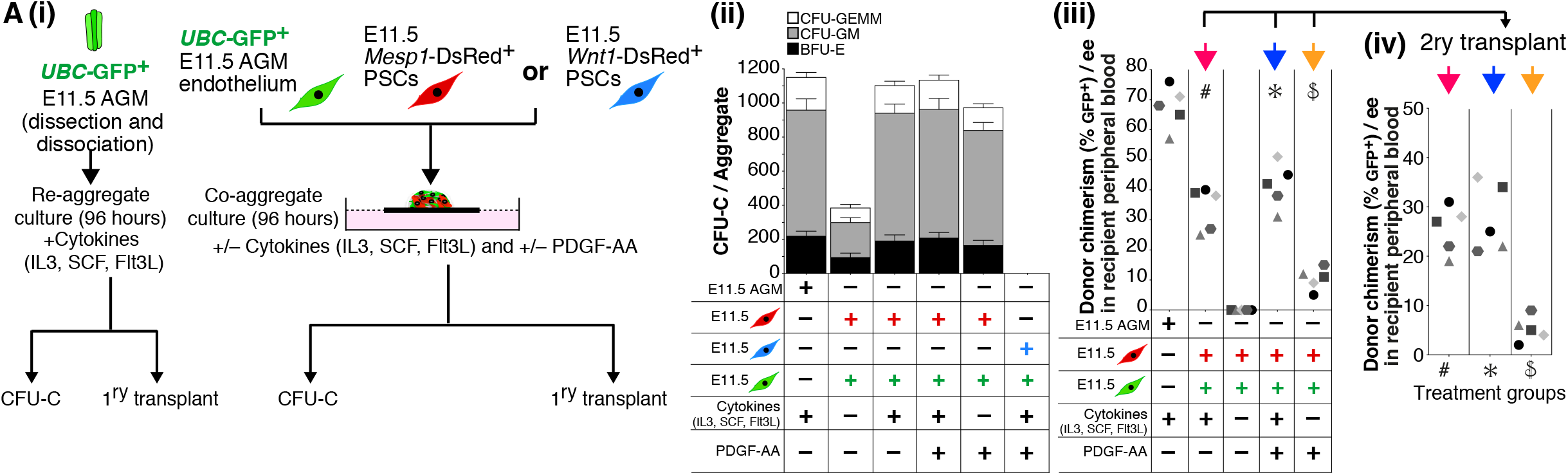
PDGF-AA can partially compensate for the absence of cytokines in co-aggregate cultures. (A) (i) Schematic diagram showing the experimental strategy for co-aggregation of E11.5 *UBC*-GFP aortic endothelial cells with *Mesp1^der^* PSCs or *Wnt1^der^* PSCs in the presence and absence of cytokines or PDGF-AA. E11.5 *UBC*-GFP AGM re-aggregates were used as controls. (ii) CFU-C assays from 96 h co-aggregate cultures (E11.5 *UBC*-GFP aortic endothelial cells/*Mesp1^der^* PSCs; n=7–10 and E11.5 *UBC*-GFP aortic endothelial cells/*Wnt1^der^* PSCs; 9–13 co-aggregates per culture condition). (iii) Percentage of GFP^+^ cells in peripheral blood in primary transplant recipients (n=5 co-aggregates per culture condition). (iv) Percentage of GFP^+^ cells in peripheral blood in secondary transplant recipients from indicated primary recipients. One co-aggregate per recipient mouse and each point represents an individual recipient. AGM: aorta gonad mesonephros, AoE; adult aortic endothelium, HE; heart endothelium, PSC; PDGFRA+ stromal cells

Movie 1: Beating cardiomyocytes from differentiated E11.5 AGM PDGFRA^+^/ *Nestin*-GFP^−^ cells.

Movie 2: Z-stack reconstruction of a matrigel plug loaded with PDGFRA^+^ *Nestin*-GFP^−^ CD31^−^ PDGFRB^−^ FAC-sorted cells from E11.5 *Nestin*-GFP^+^ AGMs and harvested at 3 weeks after transplantation.

## EXTENDED METHODS

### Mice

Mice were housed and bred in the Biological Resource Centre at the Lowy Cancer Research Centre, UNSW, Sydney. Embryos were generated by pairing respective strains of male and female mice as shown in the accompanying figures. The morning after discovery of a vaginal plug was designated as day 0.5. The developmental stage of embryos was determined according to Theiler’s criteria. All animal experiments were approved by the Animal Ethics Committee of UNSW, Sydney. Mouse strains and relevant references are listed in the Key Resource Table.

### AGM PSC isolation and *ex vivo* expansion

E10.5–E13.5 AGM regions were dissected and dissociated as previously described ^1^. E10.5–E13.5 AGM PSCs were isolated from wild-type C57BL/6 and transgenic embryos and cultured as previously described ^2^. Dissected AGMs were transferred into collagenase Type II (263 U/ml; Worthington Biosciences, Lakewood, NJ) and placed on a shaker at 37 °C for 15 min. The supernatant was passed through a 40 µm filter into a fresh tube and inactivated with 100% FCS. Cells were washed twice in 2% FCS in PBS and plated inαMEM (Invitrogen, Carlsbad, CA) with 20% FCS, and penicillin/streptomycin/glutamine (P/S/G, Invitrogen) ^2^ and cultured in the incubator at 37 °C, 5% CO_2,_ for 72 h. At the end of 72 h, cells were washed in PBS to remove non-adherent cells and cultures were continued in fresh medium. Cells were passaged on reaching 80% confluence. After passaging, cells were placed back in tissue culture flasks with αMEM + 20% FCS + P/S/G for bulk passaging. Cells were routinely cryopreserved in 10% DMSO and 90% culture medium.

### AGM PSCs’ long-term growth and serial clonogenicity

AGM PSCs were expanded in bulk culture after plating 10,000 cells per T25 flask. Resulting cells were split every 12 days. Cumulative cell numbers were calculated and plotted (log10 scale). To investigate single-cell serial clonogenicity, AGM PSC colonies were isolated individually using cloning cylinders (“O” rings) (Sigma-Aldrich). AGM PSCs micro (n=66), small (n=28), and large (n=27) colonies 2ry, 3ry, and 4ry colony formation was evaluated by plating single cells from individual colonies into 96 well plates. Cells were in αMEM (Invitrogen) with 20% FCS, and penicillin/streptomycin/glutamine (P/S/G, Invitrogen) ^2^ and cultured in the incubator at 37 °C, 5% CO_2_, for 72 h. At the end of 72 h, cells were washed in PBS to remove non-adherent cells and cultures were continued in fresh medium. Cell culture was ended at the end of day 12, and colonies were stained with crystal violet. Colony count was performed according to Chong et al. ^3^.

### AGM pericyte culture

E11.5 AGMs were dissected from *Nestin*-GFP embryos as previously described ^1^. Dissected AGMs were transferred into collagenase Type II (263 U/ml; Worthington Biosciences, Lakewood, NJ) and placed on a shaker at 37 °C for 15 min. The supernatant was passed through a 40 µm filter into a fresh tube and inactivated with 100% FCS. Cells were washed twice and cultured as described ^4^ in the incubator at 37 °C, 5% CO_2_, for 72 h. At the end of 72 h, cells were washed in PBS to remove non-adherent cells and cultures were continued in fresh medium.

### *Ex vivo* post implant embryo culture

Harvested E10.0 embryos were cultured *ex vivo* according to Sturm and Tam (1993 and 1998) protocol for 36 hours in the presence or absence of the PDGFRA inhibitor. At the end of *ex vivo* culture embryos were either fixed in 4% PFA for 15–20 min at room temperature (RT) for confocal microscopy imaging or AGM regions were dissected and dissociated as previously described ^1^ for CFU-C and CFU-F experiments. Fixed embryos were washed and embedded in OCT by flash freezing on dry ice and cut into 30 µm sections, then permeabilized with 0.03% Tween-20 in PBS (v/v) for 15 min at RT. The cells were washed once with PBS and then blocked with 10% donkey serum (v/v) in PBS for 1 h. The sections were subsequently incubated overnight at 4 °C with the primary antibodies in 2% bovine serum albumin (BSA) (w/v) in PBS, stained accordingly with secondary antibodies in 2% BSA and incubated for 1 h at 4 °C. Nuclear staining was done with DAPI. Slides were mounted with ProLong Gold mounting medium (Invitrogen). Slides were analysed using L780 LSM Zeiss confocal microscope.

### Confocal microscopy

Dissected AGM and tissues were washed with PBS (Invitrogen) for 10 min and then fixed with 4% paraformaldehyde (ProSciTech) in PBS (w/v) for 15–20 min at room temperature (RT). These tissues were embedded in optimal cutting temperature compound (OCT) by flash freezing on dry ice, cut into 30 µm sections, and permeabilized with 0.03% Tween-20 in PBS (v/v) for 15 min at RT. The cells were washed once with PBS and then blocked with 10% donkey serum (v/v) in PBS for 1 h. The sections were subsequently incubated overnight at 4 °C with the primary antibodies in 2% bovine serum albumin (BSA) (w/v) in PBS, stained accordingly with secondary antibodies in 2% BSA, and incubated for 1 h at 4 °C. Nuclear staining was done with DAPI. Slides were mounted with ProLong Gold mounting medium (Invitrogen). Slides were analysed using either a L780 LSM Zeiss or a Leica SP5 CW STED confocal microscope. Microscope and laser settings were adjusted such that no fluorescence was observed using nonimmune controls. All test specimens were observed using these same settings. 3D rendering was performed using Imaris software, to provide improved spatial information in the Z-direction. Here, we created 3D isosurface renderings from confocal Z-stacks of subcutaneously transplanted AGM PSCs in Matrigel for 3 weeks. Antibodies are itemized in the Key Resource Table.

### PDGFRA^+^ cell depletion experiments

*Pdgfra-cre^ERT^*^2^ females were time-mated with *iDTR* males to generate *Pdgfra-cre^ERT^*^2^*/iDTR* embryos. When female mice were identified as pregnant, a single injection of tamoxifen (143 mg/kg) was administered at E9.5 and after 24 hours a single injection of diphtheria toxin (4 μg/kg) was administered intraperitoneally. Embryos were harvested at E11.5 for analysis.

### *In vitro* lineage differentiation

#### Osteogenic differentiation

Osteogenic differentiation was promoted by culturing cells for 21 days in either a 6-well plate or a 4-chamber slide, containing Dulbecco’s Minimum Essential Medium-Low glucose (DMEM-LG) (Invitrogen), 10% FCS, 100 µg/ml penicillin and 250 ng/ml streptomycin, 200 mM L-Glutamine and 0.1 µM dexamethasone (Sigma-Aldrich), 10 mM β-glycerophosphate (Sigma-Aldrich), and 200 µM L-ascorbic acid 2-phosphate (Sigma-Aldrich). The cells were stained with Alizarin Red.

#### Chondrogenic differentiation

2.5 x 10^5^ cells to 1 x 10^5^ were plated in either a 6-well plate or a 4-well chamber slide, containing serum-free Dulbecco’s Minimum Essential Medium-High glucose (DMEM-HG), 100 µg/ml penicillin and 250 ng/ml streptomycin, 200 mM L-Glutamine, 50 µg/ml insulin-transferrinselenious (ITS) acid mix (BD Biosciences), 2 mM L-ascorbic acid 2-phosphate (Sigma-Aldrich), 1 mM sodium pyruvate, 0.1 µM dexamethasone (Sigma-Aldrich), and 10 ng/ml transforming growth factor β3 (TGF-β3; LSBio). Media were changed every 4 days for 28 days. Differentiated cells were stained for sulfated proteoglycans with 1% Alcian Blue.

#### Adipogenic differentiation

Cells were cultured for 7–10 days in DMEM-HG containing 10% FCS, 100 µg/ml penicillin and 250 ng/ml streptomycin, 200 mM L-Glutamine and 0.5 µM 1-methyl-3-isobutyl methylxanthine (Sigma-Aldrich), 1 µM dexamethasone (Sigma-Aldrich), 10 µg insulin (Sigma-Aldrich), 200 µM indomethacin (Sigma-Aldrich). The cells were fixed and stained with Oil Red O.

#### Smooth muscle differentiation

Smooth muscle differentiation was promoted by culturing the cells in the presence of 50 ng/ml platelet-derived growth factor BB (PDGF-BB) (R and D Systems) made up with 5% FCS in DMEM-HG, 100 µg/ml penicillin, 250 ng/ml streptomycin, and 200 mM L-Glutamine. The cells were induced for 14 days, with the medium changed every 3–4 days. The cells were stained for smooth-muscle myosin heavy chain (MYH1) and serum response factor (SRF).

#### Endothelial differentiation

Endothelial cell differentiation was promoted by culturing the cells in 5% FCS, in Iscove’s modified Dulbecco’s Medium (Invitrogen) containing 10 ng/ml bFGF and 10 ng/ml vascular endothelial growth factor (Vegf; RND Systems, 493-MV), 100 µg/ml penicillin, 250 ng/ml streptomycin, 200 mM L-Glutamine. Cells were stained for CD31. For low-density lipoprotein (LDL) uptake, acetylated apoprotein-LDL (AcLDL-Alexa Fluor 488 molecular probes) at a final concentration of 10 µg/ml was added to endothelial differentiation assays at the end of day 14. Then cells were cultured for a further 24 h. At the end of day 15, cells were fixed and uptake was assessed by fluorescence yield. For Matrigel assays, AGM MSC-LCs were plated on chamber slides containing Matrigel and cultured for 7 days. At the end of day 7, tubes were fixed and stained for CD31 and PDGFRB expression.

#### Cardiomyocyte differentiation

To promote cardiomyocyte differentiation, cells were first cultured for 4–5 days in 2% Matrigel-coated chamber slides or glass-bottom Petri dishes, in normal MSC medium. Cells were then differentiated toward cardiomyocytes in cardiomyogenic differentiation medium consisting of DMEM-LG : Medium 199 (4:1), 1.0 mg/ml bovine insulin, 0.55 mg/ml human transferrin, 0.5 µg/ml sodium selenite, 50 mg/ml bovine serum albumin, and 0.47 µg/ml linoleic acid, 10^−4^ M ascorbate phosphate, 10^−9^ M dexamethasone, 100 µg/ml penicillin and 250 ng/ml streptomycin, 200 mM L-Glutamine and 10% FCS with 1 ng/ml recombinant human neuregulin 1β2 for 14–21 days. Media were changed every 3 days. The cells were stained for cardiac α-sarcomeric actinin. Images of beating cardiomyocytes were acquired on a Nikon Ti-E microscope with a 20x phase objective (0.45 NA). 1000 frames were acquired continuously with a 52 ms frame rate. 12-bit images were acquired with a 1280×1024 pixel array.

#### Neuronal differentiation

When the cells were at 80% confluence, the culture medium was switched to DMEM-HG containing 100 µg/ml penicillin, 250 ng/ml streptomycin, 200 mM L-Glutamine, and 1 mM β-mercaptoethanol. Media were changed every 3–4 days, and culturing was for 8–10 days. Neural differentiation was confirmed by expression of *Neuron specific beta III Tubulin* (*Tuj1*).

#### Hepatocyte differentiation

At 80% cell confluence, culture medium was switched to serum-free DMEM-HG containing 100 µg/ml penicillin, 250 ng/ml streptomycin, 200 mM L-Glutamine, 20 ng/ml EGF (R and D Systems) and 10 ng/ml of bFGF (R and D Systems) to inhibit cell proliferation for 2 days. After conditioning the cells, differentiation medium was added consisting of DMEM-HG supplemented with 20 ng/ml of HGF (R & D Systems) and 10 ng/ml of bFGF, for 7 days. The cells were then cultured in DMEM-HG supplemented with 20 ng/ml OSM, 1 µmol/L dexamethasone, 10 µl/ml ITS premix, and 100 µg/ml penicillin and 250 ng/ml streptomycin for 14 days. Media were changed every 7 days. Hepatic differentiation was assessed by immunofluorescence staining for albumin (Alb) and hepatocyte nuclear factor 4 *alpha* (*Hnf4a*).

### Immuno-histochemistry

Cells were washed with PBS (Invitrogen) for 10 min. The cells were then fixed with 4% paraformaldehyde (ProSciTech) in PBS (w/v) for 15–20 min and then permeabilized with 0.03% Tween-20 in PBS (v/v) for 15 min at room temperature (RT). The cells were washed once with PBS and then blocked with 10% donkey serum (v/v) in PBS for 1 h. The cells were subsequently incubated overnight at 4 °C with the primary antibodies in 2% bovine serum albumin (BSA) (w/v) in PBS, stained accordingly with secondary antibodies in 2% BSA and incubated for 1 h at 4 °C. Nuclear staining was done with DAPI. Slides were mounted with ProLong Gold mounting medium (Invitrogen). Slides were analysed using either L780 LSM Zeiss confocal microscope or Leica SP5 CW STED confocal microscope. 3D rendering was performed using Imaris software, to give improved spatial information in Z-direction. Antibodies are listed in the Key Resource Table.

### Long-bone immuno-histochemistry

Long bones (femur and tibia) were fixed in 4% PFA for 24 h and decalcified in 14% EDTA solution for a further 24–36 h. These bones were washed and embedded in OCT by flash freezing on dry ice and cut into 10 µm sections, then permeabilized with 0.03% Tween-20 in PBS (v/v) for 15 min at RT. The cells were washed once with PBS and then blocked with 10% donkey serum (v/v) in PBS for 1 hr. The sections were subsequently incubated overnight at 4 °C with the primary antibodies in 2% bovine serum albumin (BSA) (w/v) in PBS, stained accordingly with secondary antibodies in 2% BSA and incubated for 1 h at 4 °C. Nuclear staining was done with DAPI. Slides were mounted with ProLong Gold mounting medium (Invitrogen). Slides were analysed using L780 LSM Zeiss confocal microscope.

### CFU-C assay

Colony development was performed using M3434 medium (Stem Cell Technologies) according to the manufacturer’s protocol. Colonies were scored after 7 days.

### Long-term repopulation assay

Donor tissues from *Pdgfra-cre^ERT2^/R26*eYFP embryos and neonatal long bones—or AGM tissues from *Pdgfra*-nGFP/tdTomato(ROSA26)– or re-aggregates or co-aggregates from *UBC*-GFP*, UBC*-GFP/*Mesp1*-DsRed, or *UBC*-GFP/*Wnt1*-DsRed—were isolated and transplanted via the tail vein into irradiated C57BL/6J adult recipients. Numbers of transplanted cultured cells are reported throughout the article in terms of the number present in one cultured AGM. For example, one dose is equal to 100% of the cells in one cultured AGM. Donor cells were co-injected with 20,000 wild-type bone marrow cells. Donor chimerism was determined as previously described ^5^.

### Phenotypic identification of HSCs

Cell suspensions were stained using appropriate combinations of the monoclonal antibodies anti-CD45, anti-CD31, anti-Sca1, and anti-cKIT. Donor contribution was assessed by flow cytometry using endogenous GFP and DsRed. Mice demonstrating ≥4% donor-derived chimerism (contribution to both myeloid and lymphoid lineages) after a minimum of 4 months were considered to have been reconstituted.

### Flow cytometry and cell sorting

Mononuclear staining was analysed on a Fortessa (BD Biosciences). Cell sorts were performed on a BD Influx (BD Biosciences). Antibodies are listed in the Key Resource Table. FACS data were analysed using FlowJo software (TreeStar).

### Co-aggregation assays

Co-aggregates were made by reconstituting FAC-sorted cells from one AGM equivalent, or 50,000 adult –cardiac/ –aortic/ – inferior vena cava endothelial cells with 200,000 AGM PSCs. Dissociated cells were re-suspended in 10 µl of IMDM^+^ (IMDM Invitrogen) containing 20% fetal calf serum, 4 mM L-glutamine, 50 U/ml penicillin/streptomycin, 0.1 mM mercaptoethanol, 100 ng/ml IL, 100 ng/ml SCF, and 100 ng/ml Flt3L (Peprotech). Co-aggregates were made by centrifugation in a yellow tip occluded by parafilm at 300 x g for 5 min, and cultured on top of a 0.65 µm Durapore filter (Millipore, Cat. No. DVPP02500) at the gas–liquid interface as described ^6^. Tissues were maintained in 5% CO_2_ at 37 °C in a humidified incubator.

### Inhibitor assays

To investigate the signalling pathways that determine EHT, NOTCH or BMP or WNT inhibitors were added to the co-aggregate culture media (from 0 hrs and kept for 96 hrs) and cultured for 96 hrs. At the end of 96 hrs, co-aggregates were dissociated into single cell suspensions to perform CFU-Cs and transplant assays. (See key resource table for inhibitor details). Inhibitors were reconstituted according to the manufacturer’s instructions.

### Adult –cardiac, –aortic and –inferior vena cava endothelial cell isolation

Mononuclear cells were isolated from dissected 8–12-week-old hearts, aorta and inferior vena cava via mincing of tissue before digesting in 263 U/ml Collagenase type II (Worthington) in PBS at 37 °C for 30 min, with mechanical trituration at 10 min intervals and removal of myocyte debris with a 40 µm filter. Dead cells were removed using the MACS Dead Cell Removal kit (Miltenyi Biotec) before incubation with fluorophore-conjugated antibodies (Table S2) to FAC-sort endothelial cells using BD Influx (BD Biosciences) cytometer.

### RNA sequencing, principal component analysis, and hierarchical clustering

Total RNA (100 ng) from both freshly harvested and cultured cells was isolated (phenol– chloroform separation of TRIzol LS) and purified using the Qiagen RNeasy mini kit (Qiagen, Germany) according to manufacturer’s instructions. RNA quality assessment, library preparation, and sequencing were performed by the Novogen. In brief, RNA quality was assessed using the Agilent 2100 Bioanalyzer and samples with a RIN ≥ 7.0 (average RIN = 9) were further processed. RNA libraries were prepared using the Illumina TruSeq RNA Library Preparation Kit v2, and 10 nM of cDNA was used as input for high-throughput sequencing on the Illumina HiSeqX (150 bp paired-end reads). Raw sequencing reads were filtered for adapters: reads in which more than 10% of bases were unknown, reads in which more than 50% of bases were low quality (base quality < 20). The resultant high-quality reads were aligned to the mouse genome (mm10) using the software STAR (v2.5.0b) ^7^ with standard parameters. We mapped an average of 50,727,988 reads per sample, and the average alignment rate was 94.7%. Gene expression levels were quantified using HTSeq (v0.9) ^8^. Expression levels were TMM-normalized using the software package EdgeR (v3.5) in the R statistical analysis software (v3.3.3) ^9, 10^. Genome-wide expression profiles were analysed using principal component analysis (PCA) ^11, 12^. The PCA algorithm is a dimension-reduction technique that identifies directions (called principal components) along which gene expression measures are most variant. The principal components are linear combinations of the original gene expression measures, and allow visualization of genome-wide expression profiles in two or more dimensions. Hierarchical clustering with average linkage and Euclidean distance was performed using the Partek Genomics Suite (v 6.6).

### Single-cell RNA sequencing and data analysis

Endothelial cell single-cell RNA sequencing of freshly isolated adult cardiac endothelium (0 hr) and FAC-sorted adult cardiac endothelium following co-aggregation with *MesP1*^der^ PSCs for 24, 48, 72 and 96 hrs, was performed in biological triplicates in the core facility at the Garvan Institute of Medical Research, Sydney, Australia. 10X Genomics Chromium™ Single Cell 3’ platform (v3 chemistry) was used for sequencing and Illumina HiSeq 4000 Cell Ranger (v3.1.0 was used to process raw datasets including – quality control, the extraction of gene expression matrices, and the aggregation of gene expression matrices from different sequencing runs with batch effect removal. High-quality transcriptome expression (i.e. singlets; >=5000 UMIs) profiles were extracted from 9.562 single cells. To analyze the filtered gene expression counts, we wrote custom Python 3.9 scripts, available at: https://github.com/iosonofabio/scpaper_Vashe. Briefly, the raw count matrix was converted to gene names, summing over all Ensembl IDs with the same gene name, and normalised to counts per ten thousand UMIs (cptt). We then used scanpy (https://scanpy.readthedocs.io) to log the counts, calculate overdispersed features, perform PCA and UMAP embedding (https://arxiv.org/abs/1802.03426), compute a similarity graph with 10 neighbors, and cluster with the Leiden algorithm ^13^. We then used singlet (https://singlet.readthedocs.io) to make dot plots with a threshold of 0.5 cptt and UMAP projections by cluster and logged gene expression, and cumulative distributions for the expression of selected genes within each cluster. Clusters were numbered from the highest Cdh5 expressor to the highest Runx1 expressor. Pseudotime analysis was performed using scanpy (https://scanpy.readthedocs.io/en/stable/api/scanpy.tl.dpt.html). Both branched and unbranched pseudotime were tested and yielded similar results (unbranched pseudotime was finally used for Figure 5G).

### Statistical analysis

All data are presented as either mean ± sd or mean ± sem (as indicated in figure legends). Data presented in the figures reflect multiple independent experiments, performed on different days using different mice. Unless otherwise mentioned, most of the data presented in the figure panels are based on at least three independent experiments. The significance of differences was determined using a two-tailed Student’s t-test, unless otherwise stated. p>0.05 was considered not significant; **\*** p<0.05; **\*\*** p<0.01; **\*\*\*** p<0.001. In all the figures, n refers to the number of mice. All statistical analyses were performed using GraphPad Prism software v7. Analysis of results from flow cytometry used FlowJo software (TreeStar, Oregon, US). No animals were excluded from analyses. Sample sizes were selected on the basis of previous experiments. Unless otherwise indicated, results are based on three independent experiments to guarantee reproducibility of findings.

### Data availability

The RNA sequencing dataset was submitted to the Gene Expression Omnibus database, with accession number GSE114464.

### Key Resource Table

| REAGENT or RESOURCE | SOURCE | IDENTIFIER |
| --- | --- | --- |
| **Antibodies** |  |  |
| AcLDL-Alexa Fluor 488 | Invitrogen | L23380 |
| Anti-GFP | Invitrogen | A10263 |
| Anti-GFP-Alex-488 | Invitrogen | A21311 |
| Anti-GFP-Alex-647 | Invitrogen | A31852 |
| αSarcomeric actinin | Sigma | 051M4773 |
| B220-BV421 | Biolegend | 103239 |
| Calponin | Abcam | ab46794 |
| CD146 | Biolegend | 134701 |
| CD31 | Abcam | ab28364 |
| CD31-APC | eBioscience | 17-0453-82 |
| CD31-PE | BD Biosciences | 553373 |
| CD31-PE/Cy7 | BD Biosciences | 561410 |
| CD3-Alexa Fluor 647 | Biolegend | 100209 |
| CD41- PE/Cy7 | Biolegend | 133915 |
| CD43-PE/Cy7 | Biolegend | 143210 |
| CD45 | eBioscience | 14-0451-82 |
| CD45-eFluor | eBioscience | 48-0451-82 |
| CD45-FITC | Biolegend | 103107 |
| CD4-APC/CY7 | Biolegend | 100414 |
| CD8-BV421 | Biolegend | 100737 |
| cKIT | BD Bioscience | 103101 |
| cKIT-APC | Biolegend | 105811 |
| Gr1-BV421 | Biolegend | 108433 |
| **Antibodies** |  |  |
| Hnf4α | Santa Cruz Biotechnology | sc-6556 |
| Mac1-APCCy7 | Biolegend | 101226 |
| Mouse Lineage Panel | BD Pharmigen | 557791 |
| Myh11 | Thermo Fisher Scientific | MA5-11971 |
| Nestin | Millipore | MAB353 |
| PDGFRA | BD Pharmigen | 558774 |
| PDGFRA (APA5) | Biolegend | 135901 |
| PDGFRA-APC | Biolegend | 135908 |
| PDGFRA-BV421 | Biolegend | 562774 |
| PDGFRB | Biolegend | 136002 |
| PDGFRB (APB5) | Biolegend | 136002 |
| Purified Anti-YFP | Biovision | 3991-100 |
| Sca1-PE/Cy7 | Biolegend | 108114 |
| Serum response factor | Abcam | ab53147 |
| Tuj1 | Santa Cruz Biotechnology | sc-80005 |
| VE-Cadherin | Santa Cruz Biotechnology | sc-9989 |
| VE-Cadherin-PE | Biolegend | 138105 |
| **Chemicals, Peptides, and Recombinant Proteins** |  |  |
| AG1296 | Calbiochem | 658551 |
| bFGF | R & D Systems | 3139-FB |
| EGF | R & D Systems | 2028-EG |
| Flt3L | Peprtech | 250-31L |
| HGF | R & D Systems | 2207-HG |
| IL3 | Peprtech | 213-13 |
| PDGF-AA | R & D Systems | 1055-AA |
| PDGF-BB | R & D Systems | 520-BB |
| PDGFRA (APA5) | Biolegend | 135901 |
| PDGFRB (APB5) | Biolegend | 136002 |
| SCF | Peprtech | 250-03 |
| TGF-β3 | LS-G23701 | LifeSpan |
| VEGF | 493-MV | R & D Systems |
| Matrigel Matrix | 354234 | BD Biosciences |
| WIF1 | R & D Systems | 135-WF |
| Draxin | R & D Systems | 6149-DR |
| LY450139 | Sigma aldrich | SML 1938-25MG |
| MK0752 | Santa Cruz Biotechnology | CAS471905-41-6 |
| USAG1 | R & D Systems | 9008-SD |
| **Deposited Data** |  |  |
| RNA sequencing dataset | Gene Expression Omnibus | GSE114464 |
| **Experimental Models: Organisms/Strains** |  |  |
| *C57BL/6J* | ABR Moss Vale, NSW, | *C57BL/6J* |
| *Pdgfra*<sup>tm11(EGFP)Sor</sup> | Hamilton et al., 2003 | *Pdgfra*-nGFP |
| *Gt(ROSA)26Sor*<sup>tm(EYFP)Cos</sup>/J | Srinivas et al., 2001 | R26R-eYFP |
| *Tg(CAG-DsRed\*MST)1Nagy* | Vintersten et al., 2004 | Z/Red |
| *C57BL/6Tg(UBC-GFP)30Scha/J* | Schaefer et al., 2001 | *UBC-GFP* |
| **Experimental Models: Organisms/Strains** |  |  |
| *Mesp1<sup>tm2(cre)Ysa</sup>* | Saga et al., 1999 | *Mesp1-cre* |
| *Tg(Wnt1-cre)<sup>11Rth</sup>* | Danielian et al., 1998 | *Wnt1-cre* |
| *Sox1<sup>tm1(cre)Take</sup>* | Takashima et al., 2007 | *Sox1-cre* |
| *Tg(Pdgfra-cre/ERT2)1Wdr* | Rivers et al., 2008 | *Pdgfra-cre<sup>ERT2</sup>* |
| *Tg(Nes-EGFP)33Enik* | Mignone et al., 2004 | *Nestin-GFP* |
| *Gt(ROSA)26Sor<sup>tm1(HBEGF)Awai</sup>* | Buch et al., 2005 | iDTR |
| **Software and Algorithms** |  |  |
| FlowJo | TreeStar |  |
| Partek Genomics Suite (v 6.6) |  |  |
| R statistical analysis software |  |  |

